# Tau P301L mutation promotes core 4R tauopathy fibril fold through near-surface water structuring and conformational rearrangement

**DOI:** 10.1101/2023.11.28.568818

**Authors:** Michael P. Vigers, Samuel Lobo, Saeed Najafi, Austin Dubose, Karen Tsay, Pritam Ganguly, Andrew P. Longhini, Yingying Jin, Steven K. Buratto, Kenneth S. Kosik, M. Scott Shell, Joan-Emma Shea, Songi Han

## Abstract

Tau forms toxic fibrillar aggregates in a family of neurodegenerative diseases known as tauopathies. The faithful replication of tauopathy-specific fibril structures is a critical gap for developing diagnostic and therapeutic tools. This study debuts a strategy of identifying a critical segment of tau that forms a folding motif that is characteristic of a family of tauopathies and isolating it as a standalone peptide that form seeding-competent fibrils. The 19-residue jR2R3 peptide (295-313) spanning the R2/R3 splice junction of tau, in the presence of P301L, forms seeding-competent amyloid fibrils. This tau fragment contains the hydrophobic VQIVYK hexapeptide that is part of the core of every pathological tau fibril structure solved to-date and an intramolecular counter-strand that stabilizes the strand-loop-strand (SLS) motif observed in 4R tauopathy fibrils. This study shows that P301L exhibits a duality of effects: it lowers the barrier for the peptide to adopt aggregation-prone conformations and enhances the local structuring of water around the mutation site that facilitates site-specific dewetting and in-register stacking of tau to form cross β-sheets. We solve a 3 Å cryo-EM structure of jR2R3-P301L fibrils with a pseudo 2_1_ screw symmetry in which each half of the fibril’s cross-section contains two jR2R3-P301L peptides. One chain adopts a SLS fold found in 4R tauopathies that is stabilized by a second chain wrapping around the SLS fold, reminiscent of the 3-fold and 4-fold structures observed in 4R tauopathies. These jR2R3-P301L fibrils are able to template full length tau in a prion-like fashion.

**Significance Statement:** This study presents a first step towards designing a tauopathy specific aggregation pathway by engineering a minimal tau prion building block, jR2R3, that can template and propagate distinct disease folds. We present the discovery that P301L—among the widest used mutations in cell and animal models of Alzheimer’s Disease—destabilizes an aggregation-prohibiting internal hairpin and enhances the local surface water structure that serves as an entropic hotspot to exert a hyper-localized effect in jR2R3. Our study suggests that P301L may be a more suitable mutation to include in modeling 4R tauopathies than for modelling Alzheimer’s Disease, and that mutations are powerful tools for the purpose of designing of tau prion models as therapeutic tools.

## Introduction

The microtubule associated protein tau is a pathological protein that is present in insoluble neurofibrillary tangles associated with Alzheimer’s Disease (AD), frontotemporal lobar degeneration, and many other neurodegenerative diseases, collectively referred to as tauopathies. Tau exists in solution as an intrinsically disordered protein (IDP) containing a large ensemble of disordered conformations(1). In tauopathies, tau proteins misfold and stack into insoluble amyloid fibrils, with the paired helical filaments (PHFs) in neurofibrillary tangles (NFTs) of AD being the best-known example. Recent advances in cryogenic electron microscopy (Cryo-EM) allowed the determination of tau fibril structures in multiple tauopathies. Known tauopathy structures include AD(2, 3), Pick’s Disease (PD)(4), chronic traumatic encephalopathy (CTE)(5), corticobasal degeneration (CBD) (6), progressive supranuclear palsy (PSP) (7), argyrophilic grain disease (AGD)(7), aging-related tau astrogliopathy (ARTAG)(7), globular glial tauopathy (GGT)(7), familial British dementia (FBD)(7), familial Danish dementia (FDD)(7), and limbic-predominant neuronal inclusion body 4R tauopathy(7). The region of tau observed in the core of tauopathy fibrils consists of four pseudo-repeat domains (called R1 through R4), and alternative splicing leads isoforms that contain all four repeats (4R tauopathies), and those that lack the R2 domain (3R tauopathies)(8).

The precise structure of the tau fibrils and their location in the brain tissue is diagnostic of tauopathies, but their rational generation in the laboratory is a grand challenge. The property that renders these tau fibrils potentially toxic is their “prion-like” property of cell-to-cell propagation of aggregates(9–11). A prion is defined as a misfolded protein that induces templated shape changes to functional monomers upon contact, imparting the same misfolded structure upon the monomer and extending the fibrilization. The characteristic of such a tau prion could be used to replicate tauopathy specific fibrils if a prion with a known fold can be designed. However, the defining structural property of a tau prion is not known, and shape-directed folding of tau to adopt tauopathy-specific fibril structure has not been demonstrated to date. Our hypothesis is that tau prions require a stable fibrillar construct with active growing fibril ends that serve as templating interfaces. Even so, strategies to generate a fibril structure with a target fold and the mechanism of templated aggregation are unclear. Narrowing this knowledge gap is critical for the development of therapeutic or diagnostic strategies, given that screening for small molecule drugs, antibodies, and positron emission tomography (PET) imaging agents, all require that the starting protein faithfully reproduces the fibril structure observed in the disease(12).

Post-mortem tissue from tauopathy patients is not a viable clinical target due to scarcity and patient-to-patient variation. Additionally, tauopathies develop over a period of decades, forcing researchers to rely on artificial methods to induce and accelerate pathological tau aggregation. Tools such as disease mimicking mutations or cofactors are critical to achieve prion-like propagation of pathological tau within an accelerated timeframe(9, 13, 14). The current tauopathy models for developing therapeutic or diagnostic strategies rely on various pathogenic *MAPT* mutations, such as P301L or P301S, often used in tandem with V337M or R406W(14–16). The P301 site is mutated in the majority of models of tau aggregation including mice (3xTg(17), hTau.P301S(18), JNPL3(19), PS19(20), PLB1-triple(21), TauP301L(22), among others), cell lines designed to assess seeding of aggregation(14, 23), and *in vitro* studies of tau aggregation(11). No *MAPT* mutations are linked to AD, but the mutations are routinely used as AD and other tauopathy models out of necessity, underscoring the importance of structural information about the resulting mutant fibrils in these systems. Additionally, none of the patient-derived tauopathy structures solved to date include P301L/S, nor any other exonic mutation to tau. A molecular-level understanding of the effect of this mutation on misfolding of tau is critical to justify and validate the use of mutations in designing seeding-competent tau peptide fibrils and in cellular and mouse models of tauopathies(12). Since the use of aggregation-accelerating mutations is inevitable for studies of templated aggregation of tauopathy fibrils, we focus on the consequences of the most frequently used P301L/S mutation on the energetic and structural properties of tau, in both its IDP and fibril state.

In this study we rationally select a tau fragment that, within tauopathy fibrils, adopts a core disease fold and aimed to form a similar fold in a standalone peptide. Tau is too large for its misfolding and aggregation to occur in a single step. Rather, there should be a first dominant contact and folding event of the core shape that induces the rest of the tau protein to form a disease fold. The peptide must form in register intermolecular hydrogen bonds to build the cross-β fibril structure and concurrently stabilize an intramolecular protein fold that replicates a core structure contained in a tauopathy fibril. We start with the ^306^VQIVYK^311^ segment, also known as PHF6, and include an N-terminal counter-strand that stabilizes PHF6, while keeping its hydrogen-bonding backbone moieties exposed and available to maintain an active fibril growth end. An active surface to achieve templating must also have hydrophobic hotspots to facilitate site-directed association and assembly. Once a short tau peptide is designed that successfully adopts the intended fold within seed-competent fibrils, their seeding capacity for templated aggregation of longer tau monomers makes them potential targets for therapeutic strategies. The ultimate success is for these peptide fibrils to template tauopathy structures as found in tauopathy patients.

If the main function of the P301L/S mutation is to enhance the kinetics of aggregation by lowering the aggregation barrier while minimally impacting the final structure, it is a powerful tool to model tauopathies. If this mutation biases the aggregation pathway by promoting the formation of a specific core fold found in disease fibrils, then P301L/S might be an excellent tool to narrow the pathway towards the disease fibril structures that contain this specific core. We hypothesize that the latter applies. Once the effect of the most widely used disease mutation, P301L, on modulating the formation of tauopathy-specific fibrils is understood, additional mutations can be used to enhance the intended pathway, while suppressing the spurious ones.

The P301L mutation has been reported to selectively recruit 4R tau (over 3R tau)(24, 25). In this study, we will focus on producing a 4R tauopathy prion by designing 4R tauopathy-mimicking peptide that forms a fibril core with seeding competency. The core of every 4R tauopathy fibril structure solved to date, including CBD, PSP, AGD, GGT and GPT (also called LNT), contains a strand-loop-strand (SLS) motif along the fibril axis, made of an intermolecularly hydrogen-bonded β-strand followed by a loop and another β-strand. Looking at the fibril cross-section, this SLS motif forms a ‘U-shape’ along the backbone of the peptide, encompasses the P301 residue and the hydrophobic hexapeptide motif ^306^VQIVYK^311^, PHF6. The observation that the SLS ‘U-shape’ motif is a conserved inner fold in 4R tauopathy fibrils inspired the search for a tau peptide that forms this SLS structure, once fibrilized. We selected a 19-amino-acid peptide spanning residues 295-313 of 4R tau that we henceforth refer to as jR2R3 (^295^DNIKHVPGGGSVQIVYKPV^313^) as it spans the R2/R3 splice junction of tau, includes the 301 site, the fibrilization prone PHF6 motif and an N-terminal segment that could serve as an intramolecular counter-strand. We propose that the counter-strand plays two roles: (i) orients the PHF6 segment so that its hydrogen bonding backbone moieties point along the fibril axis and (ii) stabilizes a variety of U-shape folds that form an SLS structure, with such variety represented in different 4R tauopathies (Fig. 1*A*).

**Fig 1.**
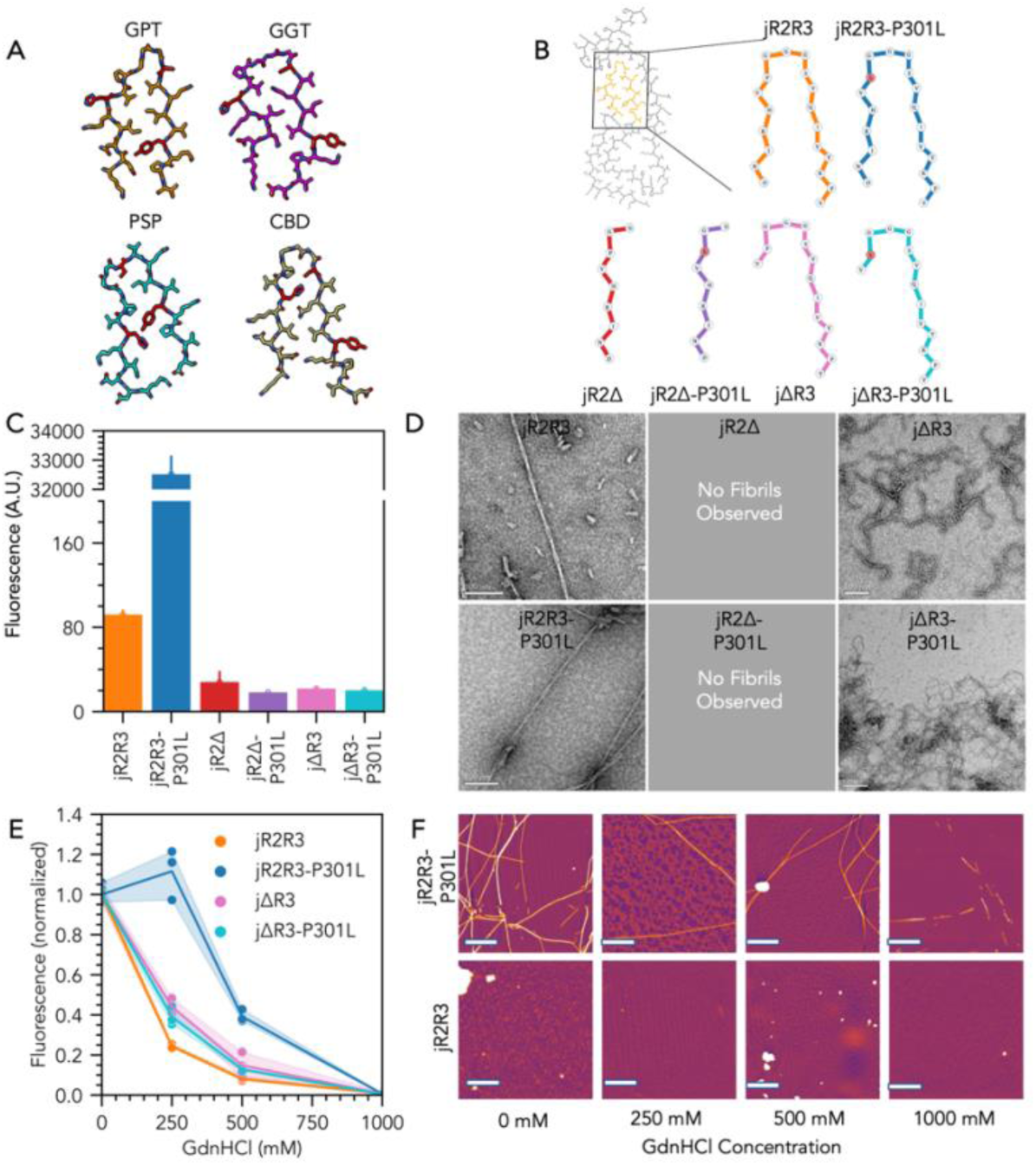
Fibril formation and stability of jR2R3 peptides: A) Structures of the jR2R3 peptide contained within known tauopathy fibril structures: GPT (PDB #: 7P6a), GGT type I (PDB #: 7P66), PSP (PDB #: 7P65), CBD type II (PDB#: 6TJX). H299, S305 and Y310 are highlight in red to help distinguish conformational differences in the SLS structures. B) Peptides used in this study shown in the conformation observed in the GPT fibril structure (PDB #: 7P6A). C) Maximum ThioflavinT fluorescence after incubation with heparin at 37°C for 18 hours. Samples contained 50µM protein with 12.5µM heparin (n=3). D) Negative stain TEM of all HP peptide fibrils formed with heparin. E) ThioflavinT fluorescence assay after incubation with guanidinium hydrochloride (GdnHCl) at 37°C for 18 hours. Samples were aggregated for 18 hours prior to denaturation. (n=3) F) AFM images of denatured samples. Fibrils were observed to degrade in a GdnHCl concentration dependent manner.

A similar tau fragment to jR2R3 was reported by Chen *et. al.* (residues 295 to 311) to contain a protective β-hairpin in solution that is opened upon mutation of P301L(26), However,, these studies did not discuss design criteria for building a tau peptide prion fibrils with the intended internal protein, which is the objective of this study.

The SLS structures are stabilized by intramolecular side chain contacts that are held together by hydrophobic and/or salt bridge interactions. In the SLS motif, the backbone is available for intermolecular H-bonding along the amyloid fibril axis. While a similar shape is formed in a β-hairpin as in a SLS strand, they are topologically distinct; a β-hairpin contains intramolecular backbone H-bonds across the hairpin that would preclude the formation of the cross-β amyloid structure(1, 26, 27). Hence, β-hairpins must be broken to populate an aggregation competent conformer to form SLS structures.

For templated aggregation to occur, breaking the aggregation-inhibiting β-hairpin to populate an aggregation competent conformer is only a first step. The interface between the active fibril-end cross section, i.e. the prion template, and the approaching tau monomer substrate must dewet. However, for the approaching tau monomer to adopt the shape of the template, a precisely coordinated process is required. The tau monomer will certainly not take up the exact prion shape prior to binding, and the dewetting of the fibril interface will not occur concurrently along the U-shaped interface all at once. Instead, we propose that in register stacking to a fibril-growing interface must be directed via a hyper-localized tau site that serves as an anchoring pin that dewets first to form the initial intermolecular contact. The characteristics of such a hyper-localized site is a small-scale (<1 nm) hydrophobic moiety surrounded by strongly water-anchoring moieties. Water then warps around this small hydrophobic moiety with a partial clathrate-like structure and greater tetrahedral ordering compared to bulk water. Such near-surface water facilitates the directed association of tau, due to an entropic gain upon eviction of the ordered, low-entropy, water at the interface, followed by a drying transition. What type of topology constitute such a site is not trivial to predict, but nonetheless locally ordered water hydrating a small hydrophobic moiety can be empirically found around folded protein surfaces(28). However, for the tau monomer to be pinned at the same hyperlocalized site, as a first step in templation, localized structuring of water has to persist in the monomer and fibril state.

Self-assembly in amyloids directed by the entropically favorable release of clathrate-like structured water has been predicted computationally (29) and structuring of water lining the active sites of enzymes has been shown to facilitate directed binding(30), but water-directed assembly to template amyloid strain has not been experimentally demonstrated. Solid-state NMR has been used in the literature to describe the hydration state of mature tau filaments(31), but the structure and thermodynamics of water solvating the tau protein in its monomeric form, prior to aggregation, are not well described. We use Overhauser Dynamic Nuclear Polarization (ODNP) to measure the equilibrium diffusivity of near-surface water around the jR2R3 peptide in its monomer state to infer hydration water structure, and state of the art computational methods to quantify the solvation free energy, structure, and dynamics of site-specific hydration water properties and thermodynamics around tau monomers in their IDP state in solution(32, 33), for the purpose of testing the hypothesis that localized water directs templated aggregation to faithfully propagate the shape of the mother fibril to the daughter fibril.

## Results

### N-terminal segment preceding the PHF6 motif is necessary to produce stable fibrils

Each of the resolved 4R tauopathy fibrils contain the jR2R3 peptide segment in a U-shaped, strand-loop-strand fibril structure. Illustrated in Figure 1*A* are the structures of jR2R3 as it exists in the known 4R tau fibril folds. Six different variants and fragments of the jR2R3 tau peptide were synthesized. They included the WT jR2R3 peptide, and the P301L mutant, designated as jR2R3-P301L. jR2Δ and jR2Δ-P301L contain the N-terminal half (295-303), whereas jΔR3 and jΔR3-P301L contain the C-terminal half (300-313) only, with the jR2Δ-P301L and jΔR3-P301L containing the P301L mutation. Each peptide variant was designed to include the 301 site as a proline (P) or leucine (L) residue. The peptide structure adopted by the 6 different jR2R3 fragments within a GPT filament are shown in Figure 1*B*.

To test aggregation capabilities, each peptide was incubated at 50µM with a 4:1 molar ratio of peptide:heparin (15kDa average MW, Galen Labs, HEP001). With the addition of heparin, all fragments containing the PHF6 segment, i.e. jR2R3, jR2R3-P301L, jΔR3 and jΔR3-P301L) formed fibrils with β-sheet content according to ThT and negative stain TEM (nsTEM) (Fig. 1*C,D*). Neither of the N-terminal half constructs (jR2Δ and jR2Δ-P301L) lacking the PHF6 motif showed any ThT fluorescence intensity, and no fibrils were observed with nsTEM, confirming that the PHF6 motif is required for fibrillization of these peptides. Because neither jR2Δ nor jR2Δ-P301L showed indications of aggregation, we conclude that the N-terminal half of jR2R3 has no inherent aggregation propensity.

Fibrils made of full length jR2R3 and jR2R3-P301L showed significant differences in β-sheet content as measured by ThT fluorescent intensity (p=2.3*10^-6^), while there was no significant difference in the total β-sheet content between the fibril made of the C-terminal half peptides, jΔR3 and jΔR3-P301L. Still, jR2R3-P301L that contains the N-terminal counter strand in addition to the PHF6 segment showed significantly greater ThT fluorescence intensity compared to jΔR3-P301L composed of the C-terminal half containing the PHF6 segment without the counter strand. The chemical nature of the leucine in comparison to proline means jR2R3-P301L can form one additional backbone hydrogen bond in amyloid fibrils, which might be a significant effect in stabilizing the resulting fibrils. However, no enhancement was observed in the quantity of jΔR3-P301L over jΔR3 fibrils (Fig. 1*C*). We can hence conclude that the effect of P301L is more complex than the addition of an extra H-bond. The pronounced enhancement of aggregation by the P301L mutation, observed only in the full jR2R3-P301L relative to the jR2R3 peptides, but not in the two C-terminal half peptides suggests that the P301L mutation alters the relationship between the aggregation-inducing C-terminal segment 300-313 and N-terminal segment 295-303. These results corroborate our hypothesis that the jR2R3-P301L fragment folds into aggregation-competent U-shape conformations stabilized into strand-loop-strand fibrils, even without the rest of the tau protein sequence.

### jR2R3-P301L fibrils display helical filament morphologies

To qualitatively evaluate the morphology and quantity of the amyloid fibrils formed, negative stain TEM was used to visualize the fibrils (Fig. 1*D*). Filamentous aggregates were observed with peptides jR2R3, jR2R3-P301L, jΔR3 and jΔR3-P301L after incubation with heparin for 18 hours. The jR2R3 and jR2R3-P301L constructs form fibril populations with distinctly different morphologies compared to that of the half peptides. jR2R3-P301L form longer and more well-defined fibrils than any other tau peptide variant. Multiple morphologies were observed within the same jR2R3-P301L fibril populations (Fig. S1), with the most prevalent one involving helical filaments with a width of 40-80 Å and a crossover length of approximately 700 Å (Fig. 1*D*). Other morphologies of jR2R3-P301L fibrils include straight, ribbon-like, or bundled filaments (**Fig. S1**). Fibrils made of jR2R3 were less abundant than jR2R3-P301L, and a straight filament morphology was observed more frequently than PHF filaments (Fig. 1*D*). The crossover distances of jR2R3-P301L fibrils are in line with those from tauopathy patient-derived brain samples, which range between 650-1000 Å. Notably, the C-terminal-only peptides jΔR3 and jΔR3-P301L formed narrow, highly tortuous fibrils with similar morphology to each other (Fig. 1*D*), corroborating our earlier finding that the P301L mutation does not influence the fibrilization path of these half-peptides in the absence of the N-terminal counter-strand.

To understand whether jR2R3 and jR2R3-P301L fibrils only differ in quantity, or also in fibril morphology and stability, we used Atomic Force Microscopy (AFM) concurrently with nsTEM. Visually, the fibril morphologies appeared similar between the TEM and AFM images, with similar fibril dimensions and periodicity observed by both techniques (Fig. S2,*A*). The height of the jR2R3 and jR2R3-P301L fibrils (Fig. S2,*B*) show that the jR2R3 and jR2R3-P301L fibrils exhibit significant differences in terms of quantity, thickness, and structural features. The jR2R3-P301L fibrils, on average, had greater heights (4-10 nm with the peak at ∼5 nm) compared to the jR2R3 fibrils (1-6 nm), indicating that fibril structures with larger cross-sections were formed. Across all samples, jR2R3-P301L also produce longer and stiffer fibrils than jR2R3, suggesting that a more stable structure is formed by *in register* packing of jR2R3-P301L to β-sheets that are less prone to fragmentation or termination.

### jR2R3-P301L fibrils are more stable than jR2R3 fibrils

To compare the stability of the jR2R3 and jR2R3-P301L fibrils, we performed a guanidinium hydrochloride (GdnHCl) denaturation assay. Fibrils were incubated with GdnHCl at concentrations ranging from 100 mM to 1 M, and ThT fluorescence measured after equilibration. A decrease in fluorescence intensity was observed in a GdnHCl concentration-dependent manner for all peptide fibrils. More stable fibrils resist denaturation under more aggressive conditions(34), so a more rapid decrease in ThT fluorescence with increasing GdnHCl concentration indicates a less stable fibril. All values were normalized to the fluorescence of the corresponding fibrils before denaturation (Fig. 1*E*). At the highest tested GdnHCl concentration (1 M), all peptides exhibited greater than 99% loss of fluorescence. The AFM images of jR2R3 and jR2R3-P301L fibrils subjected to 1 M GdnHCl treatment showed a monomer-like peptide film, indicating a complete breakdown of the filaments. The AFM of jR2R3 peptide fibrils subject to a lower GdnHCl concentration of 500 mM showed a lower total fibril density, and a reduction in fibril length and quantity instead of observing complete dissolution. At this lower GdnHCl concentration, the peptide fibrils also show clear differences between jR2R3-P301L fibrils experiencing a net loss of fluorescence intensity of 61% ± 13%, and the jR2R3 fibrils of 92% ± 1%. This implies that jR2R3-P301L produces greater quantity of more stable fibrils than jR2R3.

The fibrils of jΔR3 and jΔR3-P301L had similar stabilities to each other but were less stable than the fibrils made of the longer jR2R3-P301L peptides; all fluorescence was lost at GdnHCl concentrations greater than 250 mM. If there was a stand-alone local effect originating from the 301 site, one would expect the enhanced aggregation induced by P301L to be observed in the shorter peptides as well, but no difference in the stability between jΔR3 and jΔR3-P301L fibrils was detected. This result again indicates that the P301L mutation allows jR2R3-P301L to form more stable fibrils than jR2R3 by orienting the N-terminus as a stabilizing flanking region within the jR2R3-P301L fibrils.

### jR2R3-P301L peptides adopt U-shape fold consistent with strand-loop-strand fibril structure

We next tested the hypothesis that the jR2R3 fibrils adopt a U-fold that forms a strand-loop-strand fibril structure, given that the jR2R3 design is inspired by the different U-folds that it adopts as part of the inner core of different tauopathy fibrils (Fig. 1*A*). cryo-EM imaging with single particle analysis was performed on vitrified samples of jR2R3-P301L and jR2R3 fibrils. The jR2R3 fibrils lacked the homogeneity and helical symmetry needed to produce high-quality 2D classes from cryo-EM data, unlike jR2R3-P301L fibrils, suggesting that jR2R3-P301L adopts more well-defined folds that lead to greater stability than the jR2R3 form. jR2R3-P301L fibrils were subjected to helical reconstruction using RELION (see *SI* for more details). A 3D EM map was resolved with an estimated resolution of 3.0 Å according to the Fourier shell correlation (FSC) curve (Fig. 2*D*, Fig. S3). The resulting EM map contains a filament core with a 2_1_-screw symmetry (Fig. 2*B*). Each protofilament is composed of two jR2R3-P301L peptide chains (Fig. 2*C*). The outer chains adopts a strand-loop-strand structure, and the inner chains adopts a more extended conformation wrapping around the exterior of the strand-loop-stranded outer chain (Fig. 2*C,E*).

**Fig 2.**
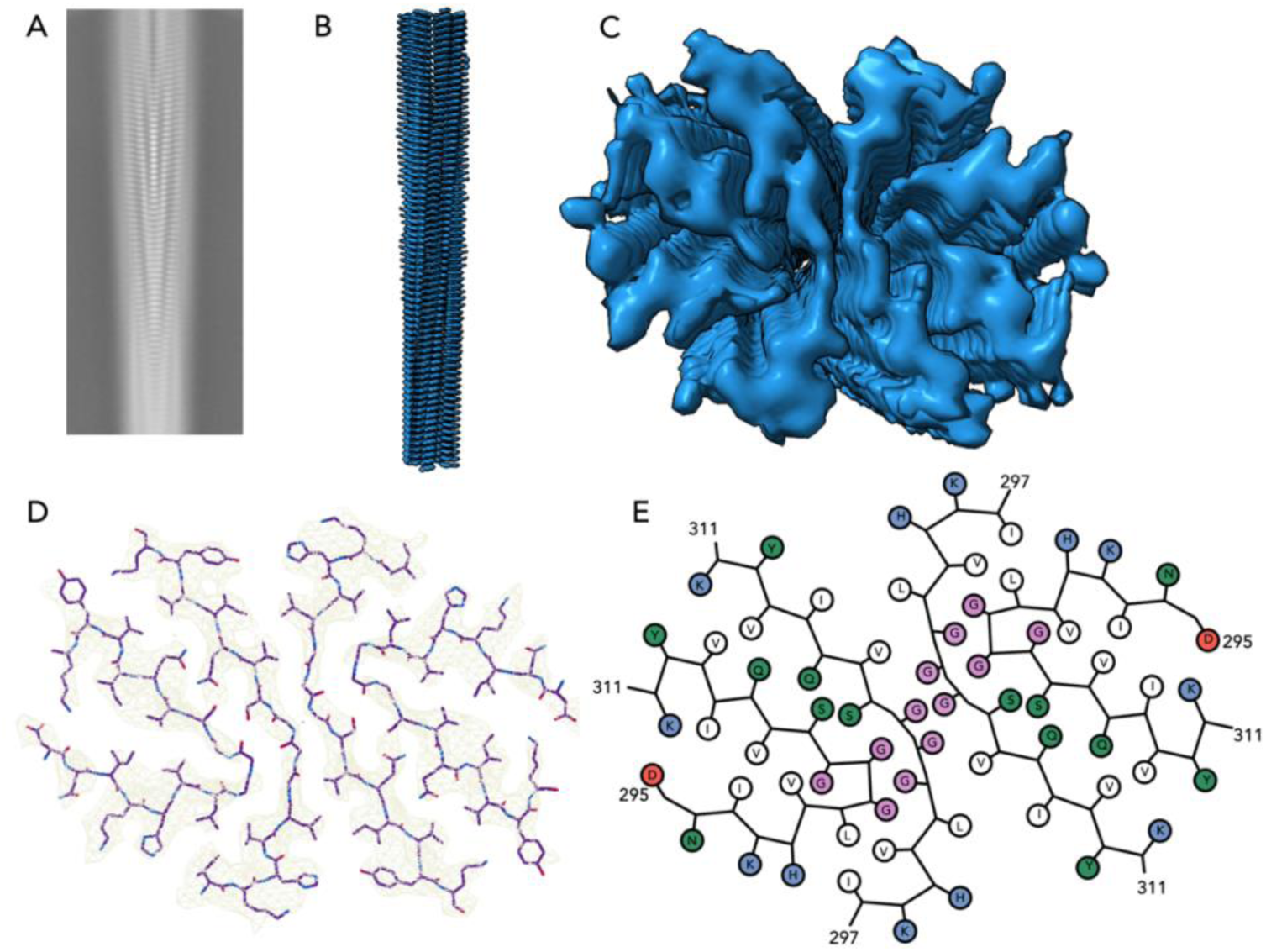
Structure of jR2R3-P301L fibrils: A) Representative 2D class average of the singlet class of fibrils. B,C) EM map of the fibril viewed from the side (B) and down the axis of the fibril (C) D) Atomic structure of a single jR2R3-P301L fibril layer (PDB ID #8V1N) E) Cartoon schematic of the jR2R3-P301L fibril layer. Pink dots are glycine residues, blue dots are positively charged, red dots negatively charged, and white are hydrophobic side chains.

The interface of the protofilaments is between ^302^GGG^304^ of both inner chains, and is stabilized by intramolecular hydrophobic interactions across sites L301 and V306. The interface between the inner (extended) and outer (SLS) chains of the protofilament are stabilized by an H-bond between site S305 of the extended chain and G303 of the SLS chain, and by a 3-residue bridge between S305 and Q307 of the SLS chain and Q307 of the extended chain. The SLS-shaped outer chains adopt a fold that is most similar to GPT in the orientation of residues ^295^DNIKHV^300^ and ^305^SVQIVY^310^. The structure of jR2R3-P301L differs from the GPT fold at site K311, which breaks the β-sheet and faces inwards the SLS fold to form a salt bridge with D295. The D295-K311 salt bridge may contribute to the closing and stabilization of the strand-loop-strand (Fig. S15*A*). The top of the strand-loop-strand might also be stabilized by P301L mutation. A backbone H-bond is formed between L301 and G304 that might help to develop the tight GGG turn that forms the SLS (Fig. S15*C*). While an exact tauopathy structure was not replicated here, the similarity with 4R tauopathy structures suggests that P301L will not preclude the replication of tauopathy structures but may inherently favor structures like CBD and GPT that have tighter GGG turns in the jR2R3 region.

In all known 4R tauopathy folds, the PHF6 region of jR2R3 is surrounded externally by the 4R repeat region of tau. In the jR2R3-P301L fibril structure, the PHF6 segment of the external U-shape fold is flanked by another extended PHF6 segment that is part of the same jR2R3-P301L peptide chain that adopts an overall extended conformation.

### P301L mutation of jR2R3 results in a greater population of strand-loop-strand fibrils

While the cryo-EM analysis revealed the structure of a subpopulation of the jR2R3-P301L fibril, it did not offer insight into the complete ensemble distribution of these structures. To measure the conformational ensemble of the tau peptides upon fibrilization, we applied double electron-electron resonance (DEER) spectroscopy to extract the probability distribution, P(r), of the intramolecular distance, *r*, between a select pair of sites of the entire jR2R3 tau peptide ensemble embedded in fibrils. DEER is a pulsed electron paramagnetic resonance (EPR) technique that probes the distribution of dipolar coupling oscillations between pairs of spin labels that are functionalized to a pair of select sites on the jR2R3 peptide prepared by cysteine mutation and spin labeling (SDSL)(35, 36). To ensure that intramolecular distances are measured within the fibrils, 10% of the peptides were doubly spin labeled and were mixed with wild type jR2R3 peptides without any cysteine mutations (see **Methods**).

To capture the difference in the fibril structure and homogeneity of jR2R3 and jR2R3-P301L folds by DEER, two sets of peptides were prepared by SDSL: jR2R3 and jR2R3-P301L peptides with spin labels attached at both ends, (sites 294 and 314), and another set with a pair of spin labels placed diagonally across the presumed U-shape fold, at the N-terminal 294 site end and another at site 305 near the center of the peptide. DEER was conducted using both sets of doubly spin labeled jR2R3 and jR2R3-P301L peptides. The end-to-end distances of the major jR2R3 and jR2R3-P301L populations in the fibril state were found to be too short to be accessed by DEER, which is consistent with the jR2R3 peptide adopting a U-shape fold in which the end spin labels are less than 1.5 nm apart—a range too short for artefact-free DEER. Further analysis with continuous wave (CW) EPR (discussed below) shows the expected proximity of the end-to-end spin labels across 294-314. In contrast, the P(r) between sites 294 and 305 for the jR2R3 and jR2R3-P301L monomers is broad as expected for an intrinsically disordered peptide, with a mean distance of 2.4 nm and spanning 2-5 nm (Figure 2). Depending on which tauopathy’s strand-loop-strand fibril structure is formed (Fig. 1*A*), the expected maximum likelihood distances between sites 294 and 305 are 2.8 nm (PSP), 3.1 nm (GGT), 3.6 nm (CBD) and 3.9 nm (GPT) (Fig. S2). In comparison to the experimental P(r) of the jR2R3 peptide monomer, the folding of jR2R3 or jR2R3-P301L into any of these folds upon fibrilization would result in a distance extension across sites 294-305. Indeed, the jR2R3-P301L(294-305) fibrils (Fig. *3B*) show a marked extension in the most probable distance across sites 294 and 305 from ∼2.4 nm in the monomer state to 3.5 nm in the fibril state, indicating that the majority population of jR2R3-P301L extends along the N-terminal half and adopt conformations closer to CBD or GPT folds.

**Fig 3.**
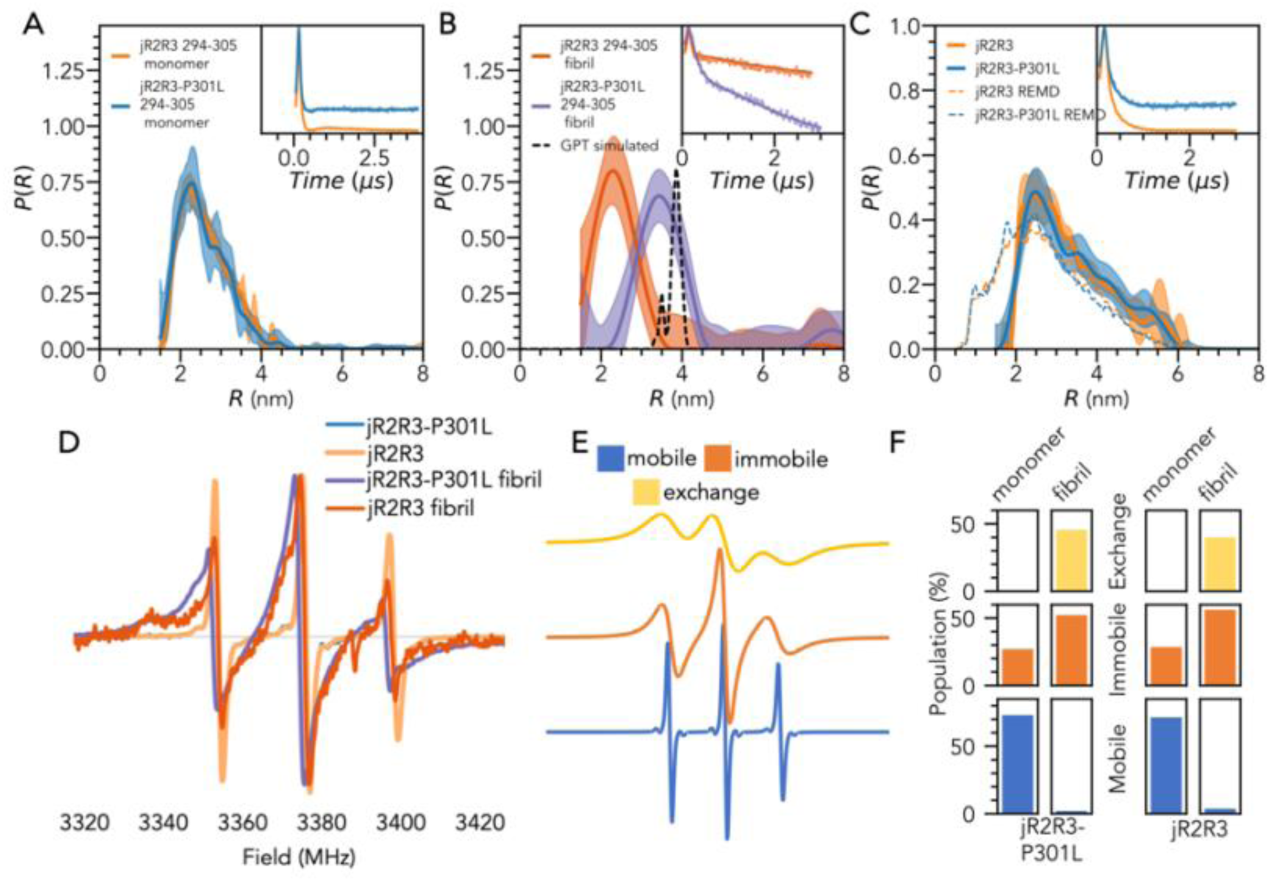
Electron paramagnetic resonance of jR2R3 and jR2R3-P301L. A) DEER probability distribution (P(r)) of jR2R3(294-305) and jR2R3-P301L(294-305) in solution state. B) P(r) of jR2R3(294-305) and jR2R3-P301L(294-305) fibers. The expected P(r) of jR2R3 in the GPT conformation is shown in dashed black. C) P(r) of jR2R3 and jR2R3-P301L labeled at sites 294 and 314 and the predicted P(r) distributions from REMD simulations (dashed line). Time domain signal is shown in inset. D) CW EPR spectra of jR2R3 and jR2R3-P301L before and after fibrilization. E) Simulated spectra of the 3 components used to fit (D). F) The proportion of the spectra attributed to mobile (blue), immobile (orange) and immobile, spin-exchanging, species (yellow).

Taken together, we surmise that the majority of the jR2R3-P301L population adopts a U-shaped conformation within the fibrils. Such distance extension was not observed for jR2R3(294-305) without P301L, upon fibril formation. These results show that the P301L mutation of the jR2R3 peptide actively facilitates the opening and folding of this peptide chain to a U-shape fold and its cross-β stacking to stable strand-loop-strand fibrils. The narrowing of the P(R) shape of fibrils made of jR2R3 labeled across sites 294 and 305 compared to the monomer state suggests that they do adopt a distinct fold(s) within fibrils and not amorphous aggregates, but the conformations of the majority population do not meet the criteria of a strand-loop-strand motif, unlike the fibrils built of jR2R3-P301L.

DEER measurements of the end-to-end distance distribution, P(*R*_ee_), of the jR2R3 and jR2R3-P301L fibrils suffered from low signal to noise ratio (SNR) and shallow modulation depth. DEER cannot detect distances below ∼1.5 nm due to spectral overlap that leads to mixing of the spin states excited by the pump and probe pulses that hence hinders the generation of quantifiable DEER modulations (37, 38). We instead acquired CW EPR spectra and performed lineshape analysis to estimate the contributions from dipolar couplings from nitroxide-nitroxide distances of below 15 Å separation. The acquired CW EPR spectra (Fig. 2*E*) were fit to a 3-component system (Fig. S4) using the MultiComponent software(39), (see Methods). The spectra were assumed to be composed of 3 major sub-populations identified in earlier studies(40): (i) a mobile component describing soluble monomers, (ii) an immobile component describing labeled peptides embedded in a fibril that is tumbling slowly, and (iii) a dipolar and/or spin-exchange broadened component describing spin-labels in a fibril within close proximity (< 15 Å) of another label (Fig. 2*G*). The CW EPR lineshape fit consisted of 86 % ± 7 % mobile species for the monomers for jR2R3 and jR2R3-P301L. In contrast, the CW EPR spectra of jR2R3 and jR2R3-P301L fibrils contained greater than 95% of the population in a fibril state, defined by either an immobile or broadened component from the 3-component fit. Of the labels incorporated into fibrils, 50% of jR2R3-P301L and 22% of jR2R3 corresponded to dipolar or spin-exchange broadened populations. With 95% of either of the peptides embedded in fibrils, but a greater proportion of jR2R3-P301L fibrils adopting a close end-to-end spin label distance compared to jR2R3, a greater fraction of jR2R3-P301L was found to adopt the U-shape fold than jR2R3. In other words, only jR2R3-P301L is forming fibrils with the majority adopting extended conformations along the PHF6 and N-terminus flank. This means that the ensemble conformation of jR2R3 peptides packed to fibrils is different from jR2R3-P301L fibrils, with a smaller population adopting U-shapes, and stacking to less well-ordered fibrils. This also answers the initial question that we posed as unknown: P301L biases the aggregation pathway by promoting the formation of a U-fold core of the SLS structure common to 4R tauopathy fibrils.

### jR2R3 fibrils can seed fibril formation of 0N4R tau

To test if jR2R3 fibrils exert prion-like seeding properties onto longer tau proteins, jR2R3 fibrils were added to 0N4R, a human full length tau variant, in their monomer form (WT and P301L) in a 1:20 molar ratio, and ThT fluorescence recorded. Both jR2R3 and jR2R3-P301L fibrils showed seeding competency in recruiting naïve full-length 0N4R tau monomers (Fig. S5). However, jR2R3-P301L fibril seeds induced aggregation of 0N4R with a shorter lag time and reached a maximum fluorescence intensity more quickly than jR2R3 fibril seeds. The ability of the 19-residue jR2R3-P301L peptide fibrils to recruit full-length tau, 20 times its length, implies that the jR2R3-P301L filament ends (either both or one) serve as a potent template to induce misfolding of intact full-length tau (of either WT or P301L) to adopt aggregation-competent conformations and states. The faster kinetics of jR2R3-P301L-seeded aggregation, in comparison to jR2R3 seeds of the same quantity, corroborates the concept that the conformation of the tau fold adopted in the strand-loop-strand fibril structure lies at the core of the templating competency of jR2R3-P301L fibrils.

### jR2R3 and jR2R3-P301L have distinct free energies despite adopting conformational ensembles with comparable end-to-end distance distributions

It has been hypothesized in the literature that WT and P301L tau adopt significantly different conformational ensembles in solution (27, 41, 42), but the associated free energy and activation barriers for adopting different conformations are still poorly understood. A previous study suggested that the proline-301 residue helps form a protective hairpin in solution of the PHF6 segment that hence inhibits aggregation of the WT form(26). The proposed hairpin was observed in molecular dynamics simulations of trimers of peptides containing residues 295-311, and the occurrence of hairpin conformations reported in the literature by cross-linking mass spectrometry. Another study detected notable local conformational differences between WT and P301L tau around V300-G303 using SAXS and NMR measurements(42).

To compare the complete ensemble of protein conformations adopted by the WT and P301L tau peptides, we used DEER to acquire the end-to-end distance distribution, P(R_ee_), of vitrified jR2R3 and jR2R3-P301L monomers in solution. The P(R_ee_) of jR2R3 and jR2R3-P301L monomers were indistinguishable from each other, with a mean R_ee_ of 26.5 Å for both (Fig. 3*C*). The populations of R_ee_ were within two standard deviations of each other at all distances. The P(R) of diagonal distances across jR2R3(294-305) and jR2R3-P301L(294-305) were similarly indistinguishable (Fig. 3*A*) within the sensitivity of DEER measurements.

Because the pairwise nature of DEER measurement does not allow for residue-level conformational analysis, molecular dynamics simulations were used to gain more insight. Reliably computing the full conformational ensemble of an IDP, such as jR2R3 and jR2R3-P301L, pushes the capabilities of current computational forcefields that tend to skew peptide conformational populations towards a more compact conformation(43). We use an integrated experimental and computational approach to first validate the computed P(R_ee_) against the experimental P(R_ee_), then select the forcefield that yields the closest agreement, and analyze the simulated conformational and energetic landscape of jR2R3 P301(L) in solution state.

The P(R_ee_) of jR2R3 and jR2R3-P301L were simulated by replica exchange molecular dynamics (REMD), a method widely used for sampling the conformational space of peptides, including tau(44–46). When using the force field a99SB-disp, recently optimized for intrinsically disordered proteins(43), the computed distributions agreed closely with the experimental P(R_ee_) distribution measured by DEER, with a computed mean end-to-end distance of 25 Å (Fig. 3*C*, Fig S6*A,B*). Next, pairwise contact frequencies of the simulated peptide ensemble were calculated, which showed a tendency for both jR2R3 and jR2R3-P301L to adopt U-shaped conformations. These U-shaped conformations (e.g. Fig. 4C*i,iii*) tend to be held by intra-molecular contacts between two regions of jR2R3: the N-terminal I297 to V300 and the C-terminal S305 to Y311 (Fig. S6*C*).

**Fig 4.**
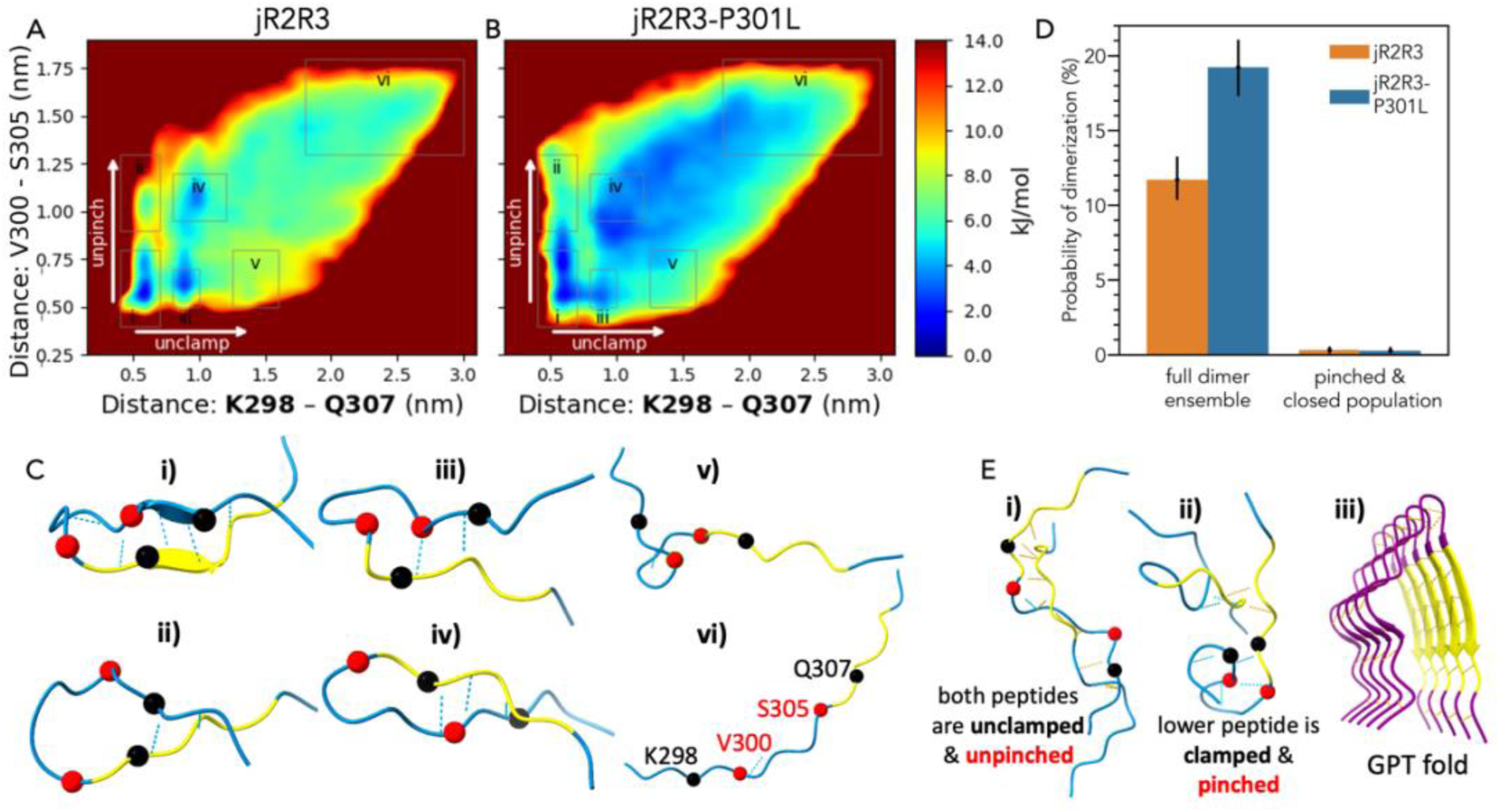
Free energy landscapes of A) jR2R3 and B) jR2R3-P301L from α-carbon distances in REMD simulations show differences in energy well depths and pathways to opening. C) Six jR2R3-P301L clusters from the various regions of the energy landscape are depicted; balls denote K298, and Q307 (black) and V300 and S305 (red), intramolecular hydrogen bonds are depicted as blue dashed lines, and the VQIVYK section is colored gold. D) jR2R3-P301L is seen to form more oligomers with 5 or more intermolecular backbone hydrogen bonds in dimer simulations than jR2R3, but these oligomer conformations rarely occur when one of the monomers is clamped & pinched, i.e. one of the monomers is in the bottom left region of the energy landscape. Error bars show 90% confidence intervals. E-i) a common cluster with many intermolecular backbone hydrogen bonds (depicted as orange lines). E-ii) A common cluster in which one monomer is clamped & pinched and only forms two intermolecular backbone hydrogen bonds. E-iii) Residues 295-313 of the CBD fold, which has many intermolecular H-bonds but no intramolecular H-bonds.

We next examined the conformational free energy landscape, which is proportional to the log of the probability distribution of distances (Fig. 4*A,B*) in order to uncover fluctuations. Infrequent fluctuations can be highly relevant to populating an aggregation pathway if they toggle between aggregation prohibiting and aggregation prone conformers. For the jR2R3 series of peptides, we focus on the free energy of breaking or forming the aggregation prohibiting, internal β-hairpin seen in the simulated contact maps. We monitored the free energy of the jR2R3 series peptides as a function of distances between two pairs of residues, across K298-Q307 and V300-S305, that form the main stabilizing contacts of the β-hairpin (Fig. 4*A-B)*. The region of the free energy landscape towards shorter distances represents conformations that are “pinched” at V300-S305 or “clamped” at K298-Q307. Here, pinching is across a short peptide region while clamping is across a broader peptide region. The bottom left of the free energy landscape (Fig. 4*C-i*) is populated with the lowest energy internal β−hairpin conformations that are confirmed to be the ones that hinder dimer formation in simulations (Fig. 4*D*). In contrast, the top right of the landscape represents open conformations (such as the ones shown in Fig. 4*C-vi*) that are less stable (i.e. higher free energy) and that we hypothesize to be aggregation competent. We found in a previous study that the region of tau around the PHF6 segment populates extended conformations in the earliest stages of aggregation when it is induced, even though the process of fibril formation, annealing and elongation may take hours, and these extended conformations may be aggregation-prone intermediates(47). These extended conformations are reported to be favored by disease mutant forms of tau(48) suggesting that suggesting the effect observed in the P301L mutation may be a more universal trait of tau pathologic mutations.

We next investigated possible transition pathways from the aggregation prohibiting β-hairpin to the open conformations. The computed free energy landscapes shown in Figs. 4A-B reveal two primary modes of unfolding towards aggregation prone conformations: unpinching at the V300-S305 contact and unclamping at the K298-Q307 contact; these modes are illustrated in the supplementary animation. The free energy landscape of jR2R3-P301L has shallower energy barriers compared to WT to break or form aggregation prohibiting internal β-hairpin conformations and allows for additional pathways to achieve unfolding that are not readily accessible to jR2R3. For jR2R3’s hairpin to open it must first unclamp (i.e. break the K298-Q307 contact) and then unpinch (i.e. break the V300-S305 contact). However, jR2R3-P301L’s hairpin can just as readily unpinch and then unclamp or unclamp then unpinch (Fig. S7). jR2R3 lacks this additional unfolding pathway due to its stiffness near residues V300 and P301, as quantified by the lower backbone dihedral entropy found only locally around these sites (Fig. 5*A*). To test the effect of backbone entropy on unpinching, jR2R3-P301L’s V300 backbone dihedrals were artificially constrained to mimic JR2R3’s V300 backbone. Under the constrained condition, the unpinch-then-unclamp mode was no longer energetically favorable even for jR2R3-P301L (Fig. S8). The free energy differences between the different conformational wells of jR2R3 and jR2R3-P301L are largely due to (**a)** the increased backbone flexibility around residues 300 and 301 in jR2R3-P301L originating from proline’s absence (Fig. 5*C*), (**b)** increased hydrogen bonding capacity of jR2R3-P301L unlocked by its more exposed conformations (Fig. S9), and (**c)** local structuring of water concurrent to increased local hydrophobicity of jR2R3-P301L (see next section) which directs tau assembly by promoting inter-molecular association selectively around these sites.

**Fig 5.**
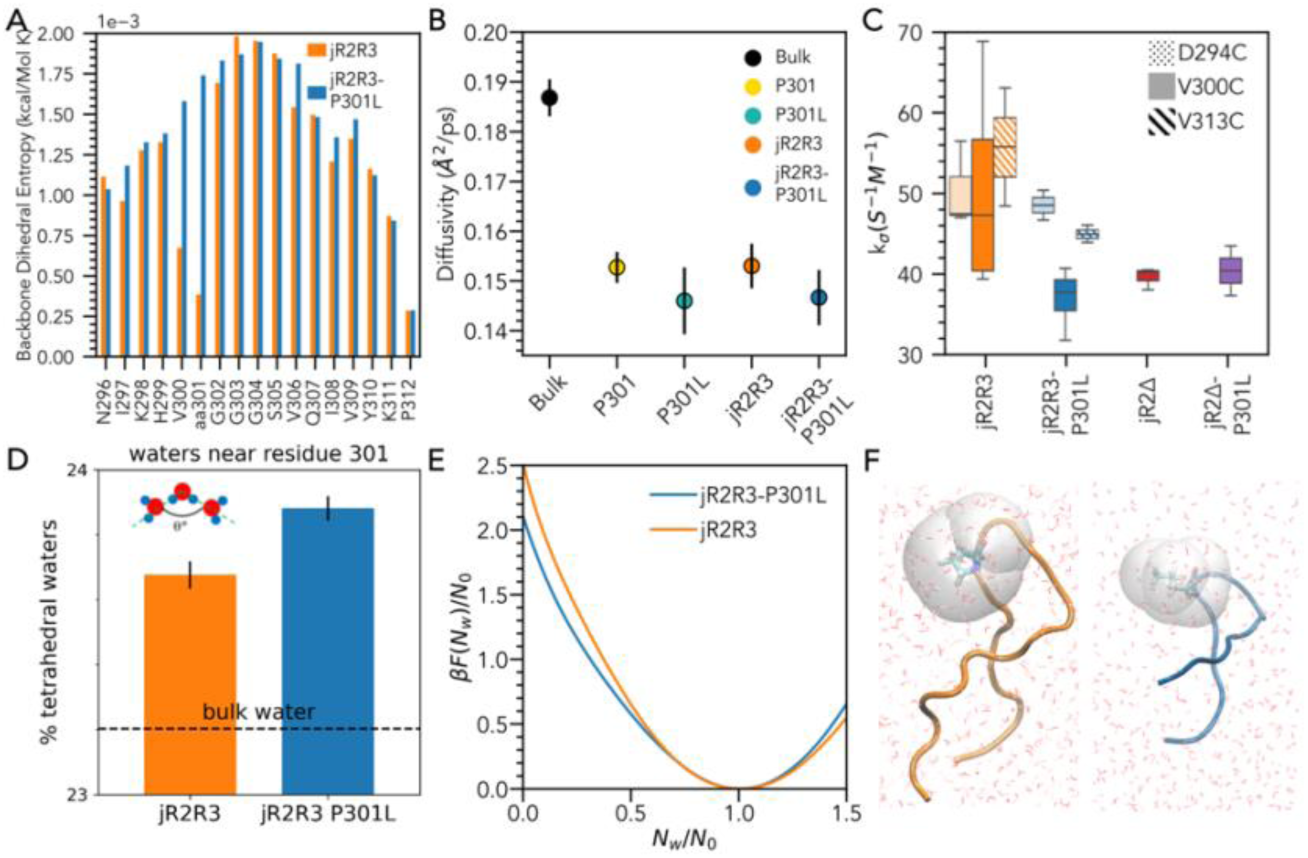
jR2R3-P301L displays altered hydration dynamics surrounding the 301 site. A) The backbone flexibility quantified by measuring the residue backbone entropy of each residue in jR2R3 and jR2R3-P301L (details in Supplement). A clear decrease in the entropy from residues V300 and P301L is seen in jR2R3. B) The water diffusivity calculation obtained from MD simulations of water directly surrounding residue 301 and on all waters surrounding the protein surface. The hydration water dynamics of the mutated peptide is both locally and globally slower than the WT peptide. C) k_σ_ calculated from ODNP measurements of jR2R3 and jR2R3-P301L was measured at three sites, the N-terminus (294C), directly adjacent to the P301(L) site (V300C), and at the C-terminus (314C) of each peptide. A significant reduction in dynamics was observed at the V300C of jR2R3-P301L in comparison the V300C of jR2R3 and 294C of jR2R3-P301L. All other comparisons were insignificant (p>0.05) with an independent t-test. n≥3 for all samples. D) jR2R3-P301L has more tetrahedral waters than jR2R3 around residue 301; this tetrahedral water fraction is associated with slower water and more hydrophobicity. Water is considered tetrahedral if its oxygen forms angles between 100°<θ<120° with adjacent water oxygens. E) The dewetting free energy per water molecule of P301 and 301L residues, N_w_ and N_0_ are the instantaneous and equilibrium number of water molecule in the probe volume, respectively, and β = 1/k_B_T, where k_B_ is the Boltzmann constant and T is the temperature. F) Representative conformations of jR2R3 (Left) and jR2R3-P301L (Right) used in umbrella sampling. Grey volume indicates probe volume from which water was expelled. ODNP also probes a similar ∼8 Å volume surrounding the tethered MTSL spin label. k_σ_ is proportional to the translational dynamics of the water of the loosely associated hydration shell.

All 4R tauopathy disease folds are held in place by a strong network of intermolecular backbone H-bonds (see orange dashes in Fig. 4*E-iii*). However, no intramolecular backbone H-bonds are found in the published structures of tau fibrils. We hypothesize that jR2R3-P301L aggregates more readily because of its shallower (around 4 kJ/mol) free energy barriers (Fig 4A-B, Fig S7) to break its aggregation-inhibiting *intra*-molecular contacts that then unlocks its ability to form ordered *inter*-molecular H-bonds that form and stabilize the fibrils. To test this hypothesis, we performed replica exchange simulations of jR2R3 and jR2R3-P301L dimers. We first validate that the most common intramolecular hairpin conformation, i.e. the pinched & clamped population prevalent in jR2R3, indeed prevents the dimer from forming intermolecular H-bonds to stabilize the fibrils (Fig. 4*D*, Fig. S10). In contrast, jR2R3-P301L readily escapes this protective pinched & clamped conformation, and can more easily form conformations that allow intermolecular H-bonds to form.

### The local water structure is perturbed near the P301L mutation site of jR2R3-P301L

While the free energy landscape of jR2R3 and jR2R3-P301L reveals that the key P301L disease mutation lowers the energetic barrier for on-pathway aggregation, it only shows that the barrier for aggregation is lowered, not whether the driving forces of association are enhanced for jR2R3-P301L. We hypothesize that unfavorable long-range interactions between like residues of jR2R3-P301L are needed to reject most association geometries that do not lead to in-register assembly to fibrils, and concurrently attractive interactions via a hyperlocalized site serve as the pinning site to align tau molecules in register. We believe that site-specific attractive interactions are mediated by structured water that hydrate small-scale (< 1 nm) hydrophobes and have enhanced tetrahedral ordering, and experience favorable interactions with other near-surface structured water with the same enhanced tetrahedral ordering. Once such sites are brought to proximity, the structured water population can be readily evicted to form a dry inter-tau interface. If the in-register aggregation of tau is to be driven by near-surface structured water that serves as pinning sites to guide and lock tau molecules to the tau fibril interface, then site-specific modulation in hydration water structure and thermodynamics in the monomer state *before* it is templated by the fibril surface is expected. Specifically, only one or two residues along the peptide should reveal signatures associated with structured water.

We hypothesize that differences in the effective hydrophobicities between jR2R3 and jR2R3-P301L -- where a hydrophobic surface is one for which the free energy of hydration, ΔG_hydration_, is positive -- contribute to the aggregation propensity of jR2R3-P301L. Previous work, including our own, showed that local water hydration dynamics measured by Overhauser dynamic nuclear polarization (ODNP) correlate with the local water hydrogen bond structure and solvation thermodynamic properties(28, 49). ODNP relies on cross relaxation of ^1^H nuclei of water molecules induced by the free electron spin of the nitroxide spin label and is sensitive to the equilibrium dynamics of water within ∼8 Å of the spin label. We report on electron-^1^H cross-relaxivity, k_σ_, derived from ODNP measurements. In the context of this study, k_σ_ is proportional to the translational diffusion dynamics of water with correlation time in the 10’s ps to 100’s ps range. Diffusion at the k_σ_ timescale corresponds to translationally diffusing hydration water bound to the protein surface with similar or stronger hydrogen bond strength compared to bulk water (28).

A spin-label was attached adjacent to residue 301 using a V300C substitution, and the P301L mutation was found to significant slow the surrounding hydration water dynamics. A k_σ_ value of 57.9 ± 32 s^-1^*M^-1^ and a kσ of 27.3 ± 13 s^-1^*M^-1^ was measured in jR2R3 and jR2R3-P301L, respectively, around the V300C site (Fig. 5*C*). These are in the range of expected k_σ_values for water on protein or peptide surfaces, and are depressed 2-3 fold compared to value for bulk water of 95.4 s^-1^*M^-1^ (28, 50). This shows that hydration water near the V300C site is significantly slowed down in jR2R3-P301L compared to jR2R3.

To understand if the hydration dynamics are globally slower across the entire jR2R3-P301L surface or only slowed near the 300 site, ODNP measurements were performed with jR2R3 and jR2R3-P301L labeled at the N- or C- termini. We found no discernable difference between any of these sites. All sites (jR2R3-294C, jR2R3-314C, jR2R3-P301L-294C, and jR2R3-P301L-314C) displayed a k_σ_ of 50 ± 5 s^-1^*M^-1^ (Fig. 5*C*). The local hydration water dynamics near the N- and C-terminal sites that are 8 and 13 amino acids away from the 301 site respectively were not altered in the jR2R3-P301L variant. In contrast, the V300C site of jR2R3-P301L was significantly slowed, indicating that the change in water dynamics is a local effect that selectively alters the water structure nearby.

The structural and dynamical properties of hydration water are modulated by the surface geometry and chemical composition of the peptide surface. To determine if the ordering of water near site 300 in jR2R3-P301L was dependent on the local peptide sequence, we performed ODNP measurements of the truncated jR2Δ-P301L and jR2Δ peptides spin labeled at the V300C site (Fig. 5*C*). No significant difference in k_σ_ was observed between either of the truncated peptides, suggesting that the conformational properties of the jR2R3 peptide must be contributing to the local structuring of water seen near site 300 in jR2R3-P301L and the difference in water dynamics between jR2R3 and jR2R3-P301L cannot be attributed to local differences in chemistry from proline to leucine mutation alone.

To compare these ODNP measurements with computational results, we compute the hydration water diffusivity via the mean-squared displacement (MSD) of water oxygens within the hydration shell of the 301 residue, and the entire jR2R3 peptide (See methods for details). The results show slower computed water diffusivity near the 301L site and averaged over the entire peptide surface of jR2R3-P301L compared to jR2R3 (Fig. 5*B*), in excellent agreement with experiments.

### Disrupted hydration shell around hydrophobic surface favors dehydration

We hypothesize that slower water near the 300 site of the jR2R3-P301L peptide indicates a lower entropy water population relative to bulk water that, upon liberation, adds an entropic driving forces for jR2R3-P301L peptide assembly that can promote protein-protein association. To test this hypothesis, we computed the free energy of dewetting of the protein surface at the 301 residue (see simulation details in SI) and the local water-protein interaction using indirect umbrella sampling (INDUS)(51, 52). We calculated the dewetting free energy for the P301(L) hydration shell, defined as the volume within 0.55 nm of any heavy atom of the residue, capturing the first two hydration layers. Due to the intrinsically disordered nature of the jR2R3-P301L tau peptides, these calculations considered multiple conformations to fully describe the ensemble hydration state. Specifically, we obtained the dewetting free energy of site 301 for the six most probable conformations of JR2R3 and jR2R3-P301L identified through Daura cluster analysis of the REMD simulations (Fig. S12). The average dewetting free energy of residue 301 in jR2R3 and jR2R3-P301L peptides was calculated by a weighted average of this representative conformational subset. The 301L site of jR2R3-P301L exhibits a lower dewetting free energy per water molecule at all levels of hydration levels compared to the 301P site of jR2R3. In particular, at N_w_=0 (where the residue is totally dewetted) the differential in free energy is 0.44 ± 0.02 k_B_T (Fig. 5*E*). The greater hydrophobicity around site 301L sensitively depends on the environment surrounding site 301, not only the nature of the 301 residue. A library of jR2R3-P301X mutants was tested for aggregation. Surprisingly, the aggregation quantity showed moderate correlation with the nominal single-site hydropathy of the 301 residue X (see supplement for more details, Fig. S14), suggesting that the local water structuring is still influenced sensitively by the side chain moiety of site 301 that hence is exerting a hyperlocalized effect.

We next mapped out the local water structure in the hydration layer around jR2R3 and jR2R3-P301L by characterizing the population of water with tetrahedral, i.e. 109.5°, angles between water oxygens all around the peptides. Tetrahedral water structure is a good measure of local hydrophobicity, because such structuring is characteristic of the hydration shell around small (<1 nm) hydrophobes (32, 33, 53). Residue 301L of jR2R3-P301L has ∼0.20% more tetrahedral hydration waters than jR2R3’s 301P (Fig. 5*D*); while seemingly small, Jiao, et. al. reported that a similar increase was associated with a ∼0.03 Å^2^/ps reduction in water diffusivity around a peptoid system(32). The extent of tetrahedral ordering of water was alternatively evaluated by the number of pentagonal and hexagonal H-bonded water rings in the hydration layer around each residue of jR2R3 and jR2R3-P301L (54). Residues 300 and 301 of jR2R3-P3011L have more pentagonal and hexagonal water rings around them than in jR2R3 (Fig. S13 *A-C*), which is likely due to the hydrophobic L301 side chain and decreased backbone rigidity in jR2R3-P301L (Fig. 5*A*). Finally, we investigated the entropy of jR2R3-P301L and jR2R3’s hydration waters across the surface of the entire peptide, and near the 301 residue. The Shannon entropy (S_θ_) of water’s 3-body angle was calculated because S_θ_ was seen by Monroe and Shell (2019) to correlate with the excess thermodynamic entropy. At ambient temperature L301 and jR2R3-P301L have a lower S_θ_ than P301 & jR2R3 respectively, which persists and becomes more pronounced at high temperatures (Fig. S13*D*). jR2R3-P301L’s hydration waters therefore have a greater entropy difference compared to bulk water (i.e. a large ΔS_hydration-bulk_), hence facilitating hydrophobic aggregation driven by dewetting that increases ΔS_hydration-bulk_.

## Discussion

This study presents the discovery and design of a tau fragment that can form fibrils with a deliberately designed tauopathy fold and potent prion-like properties. A combined experimental and computational study was conducted to describe the mechanism of action of the P301L mutation that is the single most frequently used disease mutation to model AD. To generate disease-specific fibril structures, we need simultaneous control over at least two processes: control over the precise fold of tau and the in register stacking of the tau protein, in the desired fold, into elongated fibrils. To achieve the protein fold in tau fibril structures, we identified a 19-residue peptide at the R2-R3 junction, jR2R3, that folds into a critical strand loop strand (SLS) motif. The cryo-EM structure of jR2R3-P301L resolves a SLS fold similar to those found in GPT filaments of a limbic-predominant neuronal inclusion body 4R tauopathy (LNT)(7). The SLS chain is surrounded by an additional inner chain that is reminiscent of the 3-fold and 4-fold structures of 4R tauopathies, and may be necessary to orient PHF6 within a stable fibril structure.

The jR2R3-P301L peptide fibrils show highly potent and isoform-selective seeding and templating capacity of full-length tau *in vivo*, as presented in the companion paper Longhini et. al. (55). We propose that templated seeding of much larger tau by the mini-4R-peptide fibril is initiated by pinning the tau monomer to the fibrillar surface at the same hyperlocalized site located in the core region of the active fibril end and the approaching tau monomer. We present our discovery that site 300 or 301 is such a location, but only upon P301L mutation of jR2R3-P301L. The structured, tetrahedral water wrapped around the hyper localized site of the fibril end and around the same site of the approaching tau substrate will readily associate and dewet the interface.

Our initial motivation was to understand the effect of P301L that is widely used, but not rationally justified as a tool to mimic AD and other tauopathies in cell and mouse models. This study reveals that the molecular consequences of the P301L mutation is at least two-fold. P301L lowers the energetic barrier for opening the aggregation incompetent β-hairpin conformations and make the H-bonding backbone functionalities available to form cross-β sheets. P301L also unlocks the aggregation competency of tau via hyper-localized dewetting around the 300 and 301 sites to form intermolecular hydrophobic contacts. The potency of the P301L mutation that is well-known to enhance tau aggregation of naïve full-length tau is reproduced with a 19-residue peptide, jR2R3. In other words, the effect of P301L that is utilized in tauopathy-mimicking cell and mouse models, is rooted in a localized biophysical effect of P301L that is present even in a minimal peptide motif, jR2R3. In the cryo-EM fibril structure of jR2R3-P301L solved in this study, 301L is involved in stabilizing a tight GGG turn that connects the intra-molecular counter strands via a hydrogen bond with G304. Hence, the P301L mutation enhances the ability of the jR2R3-P301L peptide to adopt a strand-loop-strand fibril structure that is at the core of all 4R tauopathy fibril structures.

P301L is a highly potent mutation to enhance the aggregation competency and seeding potency of a wide range of tau constructs including full-length tau, and hence explains why the P301L mutation is so widely used to mimic tau pathology *in vitro*, in cell, and nearly all mouse models. Our study reveals that the P301L mutation is likely more appropriate to include in pathological models of 4R tauopathies and is not rationally justified as a tool to model AD with the current knowledge available to us. This study showcases progress towards designing a peptide motif that, by design, achieves the intended fold within seeding-competent peptide fibrils. The jR2R3-P301L fibrils are expected to template the full-length tau in the same region, but more work is needed to understand if the adoption of the SLS structure around the jR2R3 region biases the folding and aggregation of the remaining region of the full-length tau. The ultimate success will be for the so-designed tau peptide fibrils to template full-length tau to adopt tauopathy structures.

## Materials and Methods

### Peptide production

Peptides were produced by Genscript to >95% purity with no additional modifications or capping beyond addition of spin labelled cysteines at the sites noted. Peptides were hydrated in 20mM ammonium acetate buffer pH7.4 to a concentration of 2 mg/mL and were immediately aliquoted and stored at −80 °C until use.

### ThT

Thioflavin T (ThT) assays for β-sheet content were conducted with 50µM protein content unless otherwise noted, and 20mM ThT in a 384-well Corning plate. A BioTek synergy2 plate reader was used with temperature control set at 37°C. Excitation used a 440 nm filter, and emission was detected at 485 nm.

### TEM

Negative stain transmission electron microscopy was performed with a 200kV FEI Tecnai G2 Sphera Microscope. 200-mesh Formvar copper grids were glow discharged for 45s with an PELCO easiGlow discharge cleaning system. 4 µL of sample was applied to grid for 1 minute, blotted with whatman paper, then 4 µL of 1% uranyl acetate was applied to the grids surface and immediately blotted. An additional 4 µL of stain was applied for 1 minute before blotting until dry.

### CryoEM

To determine the appropriate protein concentrations, buffer conditions, sample preparation and vitrification parameters, cryo-EM was performed in the MMF using G2 sphera Tecnai, FEI Titan, and FEI Talos microscopes with a Gatan 626 cryo transfer holder. Grids were plasma cleaned with an IBSS chiaro for 45 seconds. Sample vitrification was performed with a Vtirobot Mark IV. Further screening and data collection was performed at PNCC with a 200kV Arctica, and 300kV Krios. Image processing and map building was performed in RELION(56–58). Initial atomic models were built using Model Angelo(59). Initial models were corrected and edited in coot(60). Phenix was used for real-space refinement and validation(61). For further details, see supplement.

### Fibril Seeding

jR2R3 fibers were purified of excess heparin by concentration and washing in 100kDa cutoff Amicon ultra concentrator tubes. Multiple washes with DI water and concentration through the filter were preformed, and then the solution was lyophilized. Fiber mass was determined by weighing after lyophilization, and fibers were resuspended to a stock solution of 2 mg/mL (1 mM). jR2R3 or jR2R3-P301L fibrils were added to 0N4R WT and 0N4R(P301L) tau monomer at a 5% molar ratio, and ThT fluorescence intensity was measured.

### DEER

DEER signal was collected for 12-24 hours until an optimal SNR was achieved. All DEER time traces were transformed into distance distributions using DeerLab software package for Python(62). The time traces were phase corrected and truncated by 300 ns to remove possible “2+1”-artifact. One-step analysis was done using the DeerLab fit function with the following models: ex-4deer model with t1, t2, and pulse length set to experiment parameters, bg_strexp model with the stretch parameter freeze to 3 (for soluble protein) and bg_homfractal model with the fractal dimensionality set to fit between 1 to 3 (for fibril samples), and dipolarmodel using Tikhonov regularization. The uncertainty analysis was done using bootstrapping method with 100 samples. The time domain fitting results are presented both as fitting to the primary data and the distance distribution fits are presented with 95% confidence intervals. samples were corrected using spectra collected of the singly labelled species (jR2R3-(P301L)-314C) to account for the shift in geometric dimensions. The DEER experiments were performed with a pulsed Q-band Bruker E580 Elexsys spectrometer, equipped with a Bruker QT-II resonator and a 300 W TWT amplifier with an output power of 20 mW for the recorded data (Applied Systems Engineering, Model 177Ka). The temperature of the cavity was maintained at 65 K using a Bruker/ColdEdge FlexLine Cryostat (Model ER 4118HV-CF100). The bridge is equipped with an Arbitrary Wave Generator to create shaped pulses for increased sensitivity. The samples were made in D_2_O buffers with 30 % (v/v) deuterated glycerol (used as the cryoprotectant)(36). To perform an experiment, approximately 40 μL of sample was added to a 3 mm OD, 2 mm ID quartz capillary and flash frozen in liquid nitrogen to preserve sample conformations.

### CW EPR

CW EPR measurements were carried out at room temperature with a Bruker EMX X-band spectrometer operating at 9.8 GHz (EMX; Bruker Biospin, Billerica, MA) and a dielectric cavity (ER 4123D; Bruker Biospin, Billerica, MA). A sample of 4.0 μL volume was loaded into a quartz capillary tube with 0.6 mm internal diameter (CV6084; VitroCom) and sealed at one end with critoseal, and then placed in the dielectric cavity for measurements. CW EPR spectra were acquired by using 6 mW of microwave power, 0.5 gauss modulation amplitude, 200 gauss sweep width, and 10 scans of signal averaging.

### ODNP

ODNP samples were prepared with 3.5 µL in a quartz capillary tube with 0.6 mm internal diameter (CV6084; VitroCom) and sealed at one end with Critoseal. The other end was sealed with beeswax. All measurements took place under 18.0±0.2 °C with constant convective cooling to prevent sample heating. An NMR probe built in-house was used, and the sample was irradiated with microwave at the central electron hyperfine transition. NMR enhancement as a function of mw power was determined with a series of 20 mw power up to 6 W. A T_1_(*p*) enhancement curve was performed with a series of 5 inversion recovery experiments as previously described. T_1,0_ was determined in separate experiments using unlabeled peptides to be ∼2.0 s for all peptides used. Data was processed using the hydrationGUI by Tom Casey, a package for processing ODNP data using DNPLab (https://thcasey3.github.io/hanlab/hydrationGUI.html, http://dnplab.net/). Most experiments used the ‘automatic process’ function but were manually inspected after processing for aberrations in the integration windowing, or phase correction. If an error was detected (such as a failure to correctly center the NMR peak, windowing and phase correction was performed manually, and a constant 10ppm window was used.

### REMD

All replica-exchange molecular dynamics (REMD) simulations of the jR2R3 and jR2R3-P301L peptides were performed using the Gromacs package (versions 2019.6 and 2020.1)(63, 64). A 7 nm rhombic dodecahedral box was used with periodic boundary conditions. The peptides were simulated with ∼7900 TIP4P-D waters and neutralizing chlorine ions with the AMBER99SB-Disp force field(43). Monomer simulations were initialized with the peptide in an extended conformation and with histidine in the HIE protonation state, although the HID protonation state was initialized for simulations in the supplementary figure. Dimer simulations were initialized with the top two clusters (from clustering with the Daura algorithm) placed ∼2 nm apart. Monomer and dimer simulations had zwitterionic termini. REMD simulations had 60 temperatures from 300K to ∼455K. The particle mesh Ewald (PME) method with a grid-spacing of 0.12 nm was used for calculating the electrostatic interactions(65). 2 fs time-steps were used with a leap-frog integrator(66). 1.2 nm cutoff distance was used for all the nonbonded interactions. The bonds with hydrogen atoms were constrained using the SETTLE algorithm(67). The REMD systems were simulated for 400-610 ns for monomers, and 500-750 ns for dimers. See *Supplement* for details on the equilibration and how the total equilibration time was chosen.

MTSL was attached to a cysteine at residues 294 and 314 (note that jR2R3 is residues 295-313) and parameterized for a99SB-disp to compare simulation with the DEER end-to-end distance distribution. See *Supplement* for details on how MTSL was parameterized and how these MTSL-capped simulations differed from the uncapped simulations.

## Acknowledgments

The study of the role of disease mutation and selecting tauopathy-specific pathways was supported by the National Institutes of Health (NIH) under Grant Number R01AG05605. The study of the minimal prion design and seeding of tau was supported by the Tau Consortium of the Rainwater charitable fund. The ODNP study of the role of water in protein interactions, and the cryo-EM sample preparation was supported by the NIH MIRA under Grant Number R35GM136411. The ODNP study of water in protein interactions was also supported by the Deutsche Forschungsgemeinschaft (DFG, German Research Foundation) under Germany’s Excellence Strategy (EXC-2033, project no. 390677874). The W. M. Keck Foundation (www.wmkeck.org) supported the ongoing experimental and computational method developments for the tau shape propagation study.

Electron microscopy was conducted at the Microscopy and Microanalysis facility at UCSB, a part of the materials research lab (MRL). The MRL Shared Experimental Facilities are supported by the MRSEC Program of the NSF under Award No. DMR 1720256; a member of the NSF-funded Materials Research Facilities Network (www.mrfn.org).

The cryo-EM data collection portion of this research was supported by NIH grant U24GM129547 and performed at the PNCC at OHSU and accessed through EMSL (grid.436923.9), a DOE Office of Science User Facility sponsored by the Office of Biological and Environmental Research. We thank Dr. Omar Davulcu for his expert assistance with grid preparation and data collection, and the staff at PNCC for providing training and guidance with sample preparation.

We acknowledge guidance and support from Prof. Dorit Hanein and Dr. Peter Van Blerkom during and after sample preparation and data collection at the PNCC, and Prof. Niels Volkmann for image reconstruction. We also thank Prof. Dan Southworth, Dr. Eric Tse, and Dr. Gregory Merz for valuable input and advise on data acquisition and image reconstruction. We thank Bill FitzGerald for contributing to the script to quantify residue backbone entropy.

## Author Contributions

M.V., A.L., S.H. designed research. S.N., P.G., S.L., M.S.Ss., J.E.S. designed simulations. M.V., A.D., A.L., K.T., Y.J. ran experiments. M.V., K.T., S.N., P.G., S.L. analyzed data. S.H., M.S.S., J.E.S., K.K. supervised the study.

