## Supplemental Information for "Tau P301L mutation promotes core 4R tauopathy fibril fold through near-surface water structuring and conformational rearrangement"

### Supporting Information Text

### Residue hydrophobicity correlates with aggregation

Surface hydrophobicity is a complex and multidimensional parameter affected by many variables, including the local chemical and conformational landscape of the surrounding residues(1, 2). To test if there is a correlation between the hydrophobicity of the 301 site and the aggregation propensity of the jR2R3 peptide, a library of peptides of the same sequence of residues 295-313 was synthesized with 11 different mutations to site 301. A ThT aggregation assay was conducted by incubating peptides with heparin at 37˚C, as described earlier. The maximum ThT fluorescence after 18 hours of aggregation was recorded and compared to the Tanford and Nozaki hydrophobicity scale(3). A linear fit was performed, and the confidence intervals were estimated through bootstrapping (Fig. S14). The result shows a correlation between the Tanford-Nozaki hydrophobicity of the residue and the maximum ThT intensity ($r^2$=0.41, p=0.0001). We also tested other hydrophobicity scales in the literature and find similar results (Fig. S14). There are multiple driving forces for aggregation of jR2R3-P301L as has been detailed in previous sections, but the hydrophobicity of the single 301 residue appears to be also enhancing the aggregation of jR2R3 peptides. Both the WT and P301L peptides are outliers with lower and higher fluorescence intensity respectively than the fit predicts, which might be explained by differences in backbone flexibility and the conformational differences that were observed in REMD simulations. In comparing the simulated conformational ensembles of P301V and P301L, the L301 site can break its intramolecular hydrogen bonds more easily which is necessary for aggregation (Fig. S14).

### Methods

### AFM sample preparation

jR2R3 and jR2R3-P301L solution mixtures were prepared similarly as TEM samples. Peptide samples were solvated and aliquoted in 20 mM Ammonium Acetate ($NH_4Ac$) buffer with a final concentration of 2 mg/mL. The peptide samples were stored in microcentrifuge tubes at 200 µM concentration in the freezer with -20/-80 ℃. 200 µM peptide aliquots were thawed and added into 20 mM $NH_4Ac$ with heparin to a final concentration of 25/50 µM of peptide and 4:1 concentration ratio of peptide to heparin. The peptide mixtures were then incubated at 37 ℃ and shaken at 650 rpm for 23 hr.

An aliquot of 4-5uL was drop-casted on a freshly cleaved mica surface (TedPella, Redding, CA) after the incubation period. The drop-casted solution was left on mica surface for about 1 min and then the excess liquid was blotted off and dried in a desiccator overnight.

### AFM imaging

The topography and phase images were acquired by MFP-3D Atomic Force Microscopy (Asylum Research, Goleta, CA) in Tapping Mode Amplitude Modulation using high resolution silicon nitrate tip (NanoAndMore, CA), with a radius <10 nm and a cantilever with spring constant about 2 N/m and a resonant frequency around 70 kHz.

**Fibril stability assay**

Guanidinium Hydrochloride denaturation assays were conducted under the same conditions as ThT assays. Reported values were detected after signal reached equilibration at least 5 hrs after initial incubation. Fibrils were diluted 2x from initial concentration into the denaturation buffer to reach the reported GdnHCl concentration.

**REMD further details:**

**Simulations with probe**

Simulations with the MTSL probe were in an 8 nm rhombic dodecahedral box with ~12,000 TIP4P-D molecules. One sodium ion was inserted to neutralize the system. The same AMBER99SB-disp force field was used, and force field parameters for MTSL were optimized (details below). 64 replicas from 292.7 K to 455.6 K were used which resulted in average exchange rates between 20-30% for the attempted exchanges every 3 ps. These temperatures were used for MTSL simulations:
292.7, 294.5, 296.3, 298.1, 300.0, 301.9, 303.8, 305.7, 307.7, 309.7, 311.7, 313.7, 315.8, 317.9, 320.0, 322.1, 324.3, 326.5, 328.7, 330.9, 333.2, 335.5, 337.8, 340.1, 342.5, 344.9, 347.3, 349.7, 352.2, 354.7, 357.2, 359.7, 362.3, 364.9, 367.5, 370.1, 372.8, 375.5, 378.2, 380.9, 383.7, 386.5, 389.3, 392.1, 395.0, 397.9, 400.8, 403.7, 406.7, 409.7, 412.7, 415.8, 418.9, 422.0, 425.2, 428.4, 431.6, 434.9, 438.2, 441.6, 445.0, 448.5, 452.0, 455.6 K
The simulations without the probe used the same temperatures, except starting at 300.0 K instead of 292.7 K. Those 60 replicas also resulted in 20-30% average exchange rates. The MTSL simulations also constrained every bond with the LINCS(4) algorithm; the simulations without the MTSL only constrained the bonds with hydrogen atoms with the SETTLE algorithm.

**Optimization of the MTSL force field:**

To develop the force field parameters for the spin probe, we performed quantum-chemical calculations of an MTSL molecule, where the $RSO_2$ group was replaced by a terminal SH group to mimic the MTSL-cysteine disulfide bond. The structure of the MTSL molecule in vacuum was optimized in two steps: First at the Hartree-Fock (HF) level with the 6-31G* basis set(5) and next at the DFT level, using the B3LYP functional with the 6-311++G(d,p) basis set. Next, the optimized geometry was used for the Merz-Singh-Kollman (MK) population analyses using the HF level of theory and the 6-31G* basis set. The use of HF/6-31G* level of theory for deriving the electrostatic potentials (ESP) on the MTSL atoms was consistent with the development of the AMBER94(6) and the AMBER99SB*-ILDN-Q(7) force fields- based on which models the atomic partial charges for the AMBER99SB-Disp force field were primarily developed. The partial charges of the MTSL atoms were derived by fitting the ESP data using the restrained electrostatic potential (RESP) method(8, 9). The equilibrium bond-lengths and the bond-angles were determined from the DFT-optimized structure of MTSL. The parameters for the proper and the improper dihedrals were derived from the pyrroline parameters reported by Xue and Skrynnikov(10) (the authors adapted the parameters for the AMBER-based force fields from a CHARMM-based force-field, originally developed by Roux et al.(11)). All other bonded and non-bonded parameters were derived from the AMBER99SB-Disp force field by analogy, considering similar hybridizations of the atoms. The MTSL probe was attached to the β-carbon of a cysteine residue by removing the hydrogen atom attached to the terminal sulphur atom of the probe. The partial charge of the terminal sulphur atom was adjusted accordingly. All the quantum-chemical calculations were performed using the Gaussian 16 suite. The RESP fitting was carried out using the AmberTools suite (version 20)(12). The Gromacs-compatible parameter files are uploaded as Supporting Information.

**Equilibration Procedure**

The systems were energy minimized for gradient descent after being solvated. Systems were first simulated for 5 ns in NPT ensemble at 1 bar pressure and 300 K temperature. THE

MTSL simulations used a Berendsen thermostat (0.1 ps relaxation time) and Berendsen barostat (1.0 ps relaxation time) for the NPT equilibration(13). The simulations without MTSL used a v-rescale thermostat (0.5 ps time constant for temperature coupling) and the Parrinello-Rahman barostat (3 ps time constant for pressure coupling for NPT equilibration(14). Next, 5 ns long NVT simulations with the Nose-Hoover thermostat (3 ps relaxation time) were performed at 300 K temperature using the average box-size from the final 3 ns of the previous NPT step(15, 16). Then the system was copied and simulated at the various replica temperatures for 5ns without exchange. Finally, the production REMD simulation started with exchange.

The monomers with MTSL were simulated for 400 ns per replica, and the last 200 ns were used for data analysis. The monomers without MTSL were simulated for 410-610 ns per replica, of which the first 50 ns were discarded. The dimers (without MTSL) were simulated for 500-750 ns per replica. jR2R3 dimers were equilibrated for 110 ns and jR2R3-P301L were equilibrated for 80 ns of the 750 ns.

The equilibration time was decided for each measurement by using a heuristic that maximizes the number of effectively uncorrelated measurements. The autocorrelation function was integrated for equilibration times in 10 ns increments starting from 50 ns, and the equilibration time with the most effectively uncorrelated measurements was chosen. Dimer simulation equilibration times were chosen by maximizing the number of effectively uncorrelated intermolecular H-bond counts. Autocorrelation functions were measured using pymbar’s timeseries module(17).

**INDUS**

The top 6 clusters from a Daura clustering analysis of the REMD monomer simulations were further analyzed with INDUS. GROMACS (version 4.5.3) was modified to bias the coarse-grained number of waters, $N_v$, in the hydration volume of the P301 or L301 residue using the Indirect Umbrella Sampling (INDUS) method(18, 19). The Gaussian coarse-graining function employed in INDUS is parameterized with a standard deviation of $\sigma$ = 0.01 nm and a truncation length $r_c$ = 0.02 nm. The water dynamics is governed by the Hamiltonian: $H = H_0 + 1/2\ \kappa\ (N_v - N^*)^2$, where $H_0$ is the unbiased Hamiltonian, and the second term represents the harmonic biasing potential with strength $\kappa$ = 10 $k_BT$. The overlap in the $N_v$ values that are obtained from sampling over the different $N^*$ water molecules (i.e. “for $N^*$ in {-6..3..54}”), enable us to estimate the dehydration free energy of the target residue for the $N_v$ values, as characterized by the logarithm of the unbiased probability distribution $P_v(N)$, of observing N water molecules in the hydration volume of the target residue that is calculated using the Unbinned Weighted Histogram Analysis Method (UWHAM)(19, 20).

The rectangular simulation box with size $L_x = L_y = L_z$ = 7.12 nm that contains a peptide and ~11800 water molecules were simulated by using the leapfrog integrator(21) to integrate the equations of motion with a time-step of 2 fs. The oxygen-hydrogen bonds in water were constrained using the SETTLE algorithm(22), and all the peptide bonds involving hydrogen atoms were constrained using the LINCS algorithm(4). The short-range van der Waals and Coulombic interactions were truncated using a cut-off of 1.0 nm, and long-range electrostatics were calculated using the particle-mesh Ewald (PME) algorithm(23). After solvating the 6 most probable conformations obtained from the REMD simulations for each jR2R3 and jR2R3-P301L peptides, the systems were energy minimized through the steepest descent approach. The following MD equilibration simulations were performed prior to the production phase. First, the systems were simulated in an NVT ensemble at temperature T = 300 K for 500 ps, by using the stochastic velocity-rescale thermostat with a time constant of 0.5 ps(24). Next, a 1.0 ns NPT simulation was performed for every system by using the Berendsen barostat(13) to maintain the temperature and pressure at 1bar. Following this NPT equilibration, we performed a 3.0 ns NPT run for every system by using the stochastic velocity-rescale thermostat with a time constant of 0.5 ps, and with employing the Parrinello-Rahman barostat(14) with a time constant of 1 ps, to maintain the temperature and pressure at 300 K and 1 bar. All the production runs were performed for 10 ns at 300 K and 1 bar, by using the stochastic velocity-rescale thermostat and the Parrinello-Rahman barostat; the first 2 ns were discarded to reach the convergence phase.

**Water Structure Analysis**

The "tetrahedral water" fraction for each residue, was calculated by integrating the water triplet angle distribution from 100-120° for each residue. The water triplet angle distribution is the distribution of angles formed by a central water oxygen and the oxygens adjacent waters. The water triplet angle distribution has been shown to reveal signatures of water structuring in response to different types of solutes and interfaces(25, 26). The distribution of hydration waters 4.25 Å around each residue was computed; the 4.25 Å cutoff corresponds to the second minimum in the RDF between the heavy atoms and water oxygen atoms. Neighboring waters are defined as those whose oxygens lie within 3.4 Å of a central water. We computed these water angles for each frame of the converged trajectory, which was saved every 10 ps, and we created a histogram of the angles with 5° bins. Each frame has statistically independent water triplet angles measurements based on measured autocorrelation times. All confidence bars were determined by bootstrapping independent measurements 1000 or more times with replacement. The Shannon entropy of this distribution was calculated by summing the p*log(p) of each of the 5° bins.

Water rings were counted using code from this repository: https://github.com/vitroid/cycless.git. We look at waters 4.25 Å around each residue and determine if two waters are connected based on whether the O-H distance between two waters is closer than 2.45 Å. We count pentagonal or hexagonal water rings when five or six waters in the residue's hydration layer form a closed, connected graph.

**Residue Backbone Entropy**

Residue backbone entropy was measured by analyzing the regions of Ramachandran space that each residue occupied. The Φ and Ψ dihedral space was coarse-grained into four quadrants as follows; A: {Φ >0°}, B : {Φ<0 , (−120°<Ψ<50°)}, C : {(−100°<Φ<0°), (Ψ>50° or Ψ<−120°)}, D: {(−180°<Φ<−100°), (Ψ>50° or Ψ<−120°)}.

Each residue in each frame is assigned a region. Finally, the residue backbone entropy ($S_{res}$) is measured by calculating the Shannon entropy of the Ramachandran regions occupied by each residue,

$$\frac{S_{res}}{k_b} = -\sum_i p_i \cdot \ln(p_i) \qquad (1)$$

where $i$ is the region of Ramachandran space and $p_i$ is the probability that the residue will be in that region.

**Other Analysis of MD Simulations**

Energy landscapes were generated by measuring the 2D probability density, taking a logarithm and scaling by Boltzmann's constant and temperature, and applying gaussian interpolation to smooth the landscape. Residues were assigned secondary structure with the DSSP algorithm(27).The Daura algorithm was used to cluster the most probably conformations(28); a 0.3 nm root-mean-squared deviations cutoff was used for the backbone $C_\alpha$-C-N atoms.

All error bars for MD analyses are 90% confidence intervals, unless otherwise stated. We used pymbar's timeseries module to calculate the statistical inefficiency. Independent measurements were bootstrapped 1000 times *with replacement* and sorted to determine the confidence levels. ChimeraX and Pymol were used to visualize conformations. MDAnalysis and mdtraj were used for data analysis. The unzipping and unpinching mode animations were created by visualizing one of the first principal components of the α-carbons coordinates for a clamped and pinched subset of the jR2R3-P301L trajectory, respectively.

**DEER P(r) simulations**

The expected P(r) distributions of the jR2R3 in the tauopathy fibril cores (Fig. 3*B*, S2*C*) were predicted using the DEER-PREdict package(29). A 298K MTSL library was used to simulate all possible conformations of the labels once tethered to the defined structure extracted from. To form the protein backbone all residues except 294-314 were deleted from each PDB structure. Three chains (A, E, C) were retained, and the labels were attached to the desired label sites of the middle chain (A) to account for clashes with adjacent chains.

**Spin-labeling**

Peptides were functionalized with an a nitroxide free-radical, MTSL (S-(1-oxyl-2,2,5,5-tetramethyl-2,5-dihydro-1H-pyrrol-3-yl) methyl methanesulfonothioate). MTSL was incubated at 10x molar concentration with peptides in 4M GdnHCl overnight at 4˚C. Labeled protein was passaged through two GE PD G-10 desalting columns and exchanged into 20 mM ammonium acetate buffer pH 7.0, with no salt present to prevent preliminary aggregation.

Peptides used to measure DEER of fibrils were labelled by mixing the DEER double mutant with 20x concentration of WT protein. MTSL was added in 10x molar excess and the solution was incubated at 4 ˚C for 1hr. Next, heparin was added in a 4:1 molar ratio (tau:heparin) and the solution was incubated at 37 ˚C for 48 hrs. To remove unbound spin-labels, fibrils were concentrated using a 50 kDa cutoff Amicon ultra concentrator, then diluted with buffer and concentrated again. This was repeated until at least 100x dilution of the original buffer was achieved.

**CWEPR spin counting**

Samples were aliquoted and frozen until use at ~100uM protein concentration. Spin-labelling efficiency was measured through quantitative cwEPR. A standard ladder of of 4-HydroxyTEMPO was measured at 50, 100, 200, 500 µM concentration and cwEPR spectra were recorded. The maximum of the baseline-corrected, integrated spectra was recorded, and fit to a linear regression vs. concentration. All samples were measured under the same conditions, and the max of the baseline-corrected, integral spectra was converted to an effective spin label concentration. Spin labelling efficiency was calculated by $SL_{eff}$=[SL]/[Protein].

**Multi-Component CWEPR spectra Fitting**

Fitting of cwEPR was performed using MultiComponent, a software package from Christian Altenbach. *The program is written in LabVIEW (National Instruments) and can be freely downloaded from the following site:* http://www.chemistry.ucla.edu/directory/hubbell-wayne-l. Fitting was done using similar procedures as described previously. Briefly, the mobile component was determined by fitting the A, G and R tensors to the solution peptides. The A and G tensors were determined first, and then R was allowed to vary. Next a second component was fit to fibrils formed with 10% spin-dilution and a single labelled site at V300C. These fibrils should display minimal exchange-coupling but exist in the slow-motion regime due to the size of the fibril. Fibrils were previously determined have an axially symmetric diffusion tensor ($\alpha_D$ = 0˚, $\beta_D$ = 36˚, $\gamma_D$ = 0˚) and an ordering parameter of 20. Only the rotational correlation time (R) was fit to this component. Finally, a 3rd component was added to fit doubly-labeled peptides in fibrils. Component 3 used the same rotational correlation time (R) as the second component, as all spin labels in the fibrils should be in similar rotational correlation regimes. A Heisenberg spin-exchange component ($\omega_{ss}$=140MHz) previously empirically determined for tau fibrils was added to component 3. Solution state peptides were fit to only the mobile (component 1) and immobile (component 2) components. When fit with a 3rd, spin-exchanged, component the covariance of component 2 and component 3 was too large to obtain trustworthy fits.

**DEER pulse program**

The following 4-pulsed DEER sequence was applied to all samples: $\pi_{obs}/2 - \tau_1 - \pi_{obs} - (t{-}\pi_{pump}) - (\tau_2{-}t) - \pi_{obs} - \tau_2$ – echo. $V(t)$ is recorded as the integral of the refocused echo as a function of time delay, $t$, between the Hahn echo and pump pulse. Rectangular observe pulses and chirp pump pulse were used with the following pulse durations: $\pi_{obs}/2$= 12 ns, $\pi_{obs}$= 24 ns, $\pi pump$= 100 ns. The chirp pump pulse was applied with a frequency width of 60 MHz to excite a distinct spin population, referred to as B spins, while the observe spulse was set 33 G up field from the center of the pump frequency range to probe another distinct spin population, A spins. $\tau_1$ was set to 180 ns and $\tau_2$ was set according to the SNR profile of the dipolar signal. Deuterium ESEEM suppression was done by incrementing $\tau_1$ with 16 ns steps, n=8 times. The data was acquired with resolution of 16 ns, 16-step phase cycling, and signal averaged until desirable SNR.

**Cryo-EM**

Multiple sample concentrations were tested, and 10 µM protein samples, in a 20 mM ammonium acetate buffer (pH 7.4) with 10 mM NaCl were determined to provide adequate sample coverage and ice thickness for data collection. Samples of jR2R3 and jR2R3-P301L fibrils were incubated at 37˚C on an Eppendorf thermomixer at 600 rpm for 18 hrs. Quantifoil 2/2 grids were prepared with either one application of 5 μL, or 2 applications of 3 μL. A single blot of 2 s was applied with a Vitrobot, in the case of 2 applications, a manual blot was applied between the first and second sample application by touching the torn edge of a filter paper to the backside of the grid. Grids were sent to PNCC for further screening and data collection. Grids were screened with a 200kV Arctica microscope. Full data collection was conducted a Titan Krios microscope with a falcon 3 detector, and bioquantum energy filter. SerialEM was used to control image acquisition(30). Fibrils were observed to partition towards the edge of holes, so 5 micrographs were collected per 2 μm hole, focused around the edge of the holes. Micrographs had an electron exposure of 50 $e^-/Å^2$, and a defocus range from -1.0 μm to -2.0 μm. Micrographs were motion corrected using the Motioncor2 implementation in RELION-4.0 (31, 32). CTF correction was performed with CTFFind4(33). Fibrils were manually picked in RELION. 1024 pixel particles were extracted 3 asymmetrical units apart and downscaled to 256 pixels. 2D classification was performed and two distinct fibrils populations emerged: a singlet (Figure 2A), and doublet fibril (Figure S3A). The doublet fibril did not contain enough particles to proceed to 3D classification or refinement. Eight 2D classes of the singlet type were used to create an 2D initial model that was used in 3D refinement. Particles were reextracted with a 386 pixel box size for 3D refinement. Particles went through multiple rounds of 3D refinement followed by CTF correction and particle polishing until a final resolution of 3.0 Å was resolved as determined by the Fourier Shell Correlation (FSC=0.143) (Figure S3B). After initial 3D refinement, a $C_2$ symmetry and a $2_1$ screw pseudo symmetry were tested, and the $2_1$ symmetry was determined to provide a better refinement of the structure and was imposed for future refinements.

Initial model building was conducted in ModelAngelo(34). The handedness of the map was inverted and provided a better fit to the jR2R3 peptide. Initial models were corrected and refined manually in coot(35). 5 layers of the fibril were refined in Phenix(36), and the model validation was conducted in phenix.

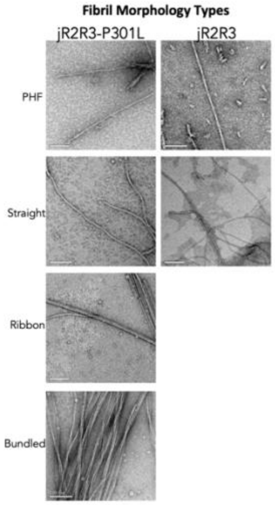


**Fig. S1.** The variety of morphologies observed in jR2R3 and jR2R3-P301L fibrils populations.

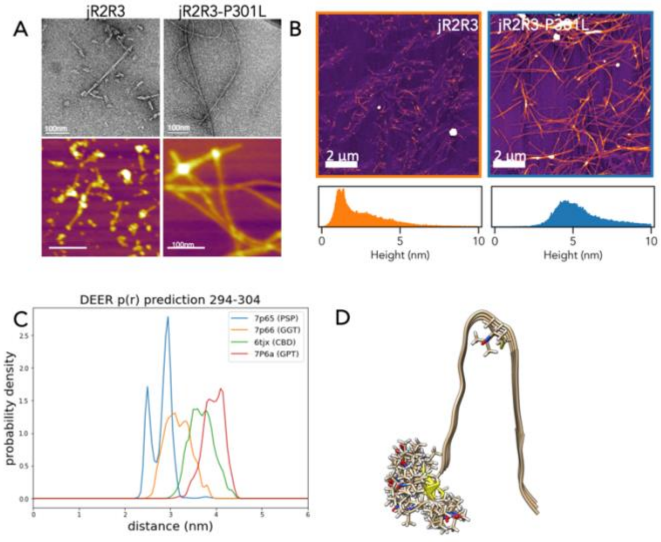


**Fig. S2**. A) Demonstration of the consistency of morphologies observed with NS-TEM and AFM. B) AFM images of jR2R3 and jR2R3-P301L fibrils (top). The height of the fibrils recorded by AFM were quantified and are plotted as histograms (bottom panel) C) Expected P(r) distance distributions between sites 294 and 304 for the 4 strand-loop-strand motifs of jR2R4 that have been reported in the literature. Note that DEER measurements shown in Figure 3A,B were measured between sites 294 and 304. Site 305 is not able to be simulated with the published structures as the sidechain is directed inward in the fold and placing a nitroxide radical leads to steric clashes. D) The structure of the strand-loop-strand motif in the 4 structures simulated in (C).

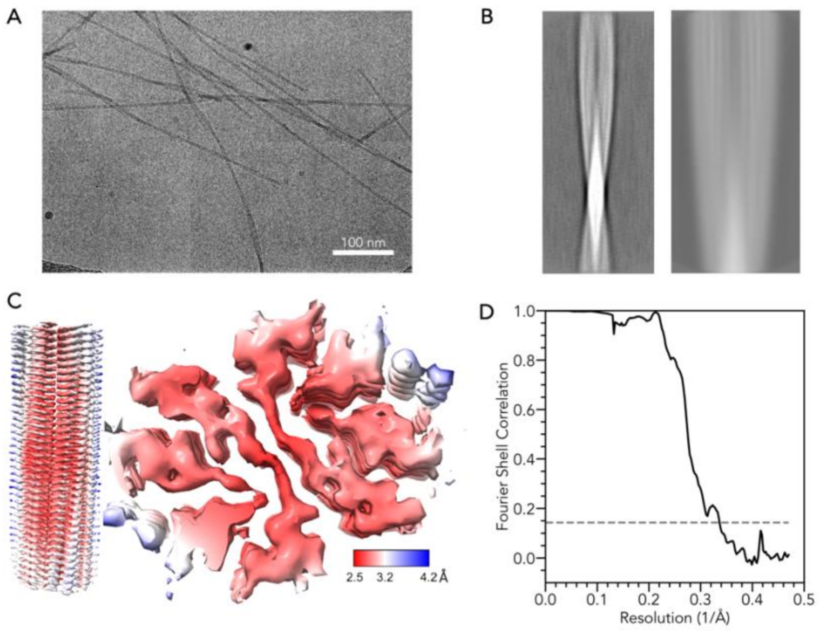


**Fig. S3.** A) Representative cryo-EM image of jR2R3-P301L fibrils. B) Example 2D class averages of the 'doublet' fibrils of jR2R3-P301L observed with CryoEM. Left, a class average from 2D classification of 1024 pixel box size particles, downscaled to 256 pixels. Right, a class average from 2D classification of 384 pixel box size particles. C) Local resolution estimation of the jR2R3-P301L EM map. (D) Gold standard FSC curve of the EM map shown in Figure 2C,D. 2D classification, resolution estimation and local resolution estimation was performed in RELION.

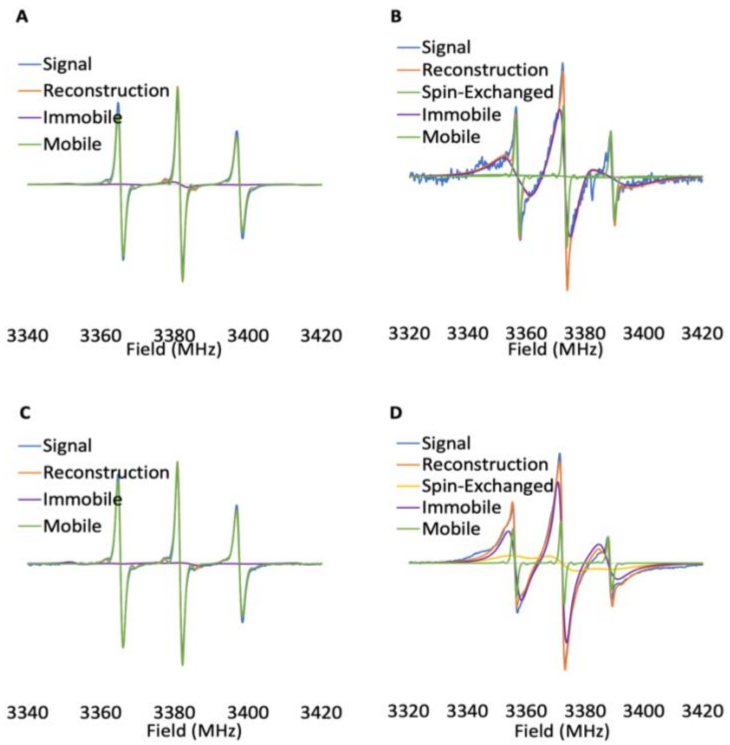


**Fig. S4.** EPR spectra (blue) fit to multiple spin components, the resulting fit (orange), and the contribution by each component (purple, green, yellow) A) jR2R3 B) jR2R3 fibers C) jR2R3-P301L D) jR2R3-P301L fibers.

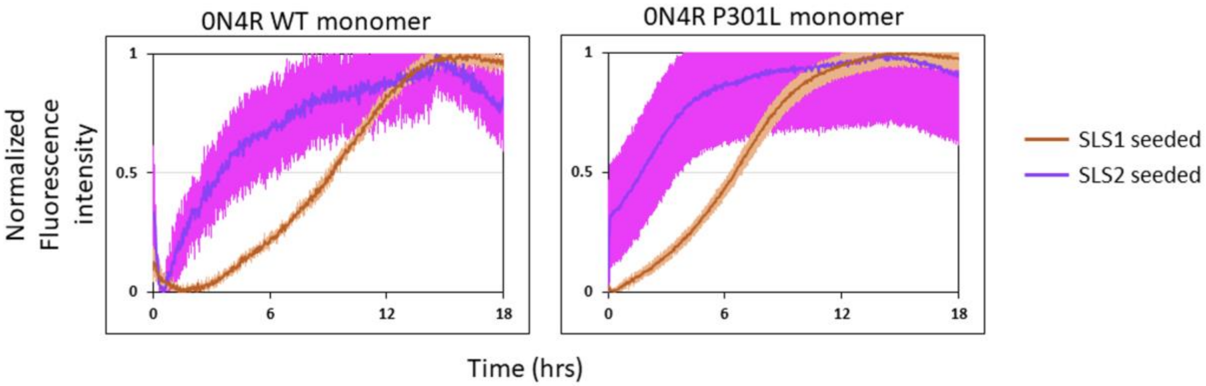


**Fig. S5:** ThT fluorescence of 0N4R WT (left) and 0N4R P301L (right) seeded by jR2R3 (red) and jR2R3-P301L (purple) fibrils.

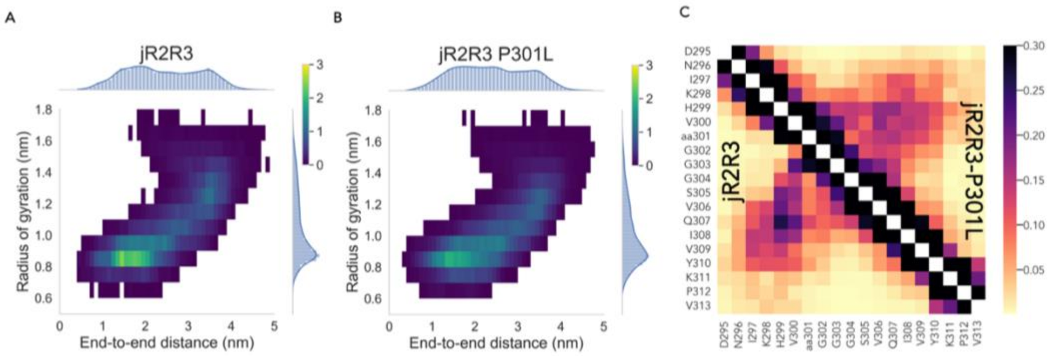


**Fig. S6**. A,B) Heat map of the end-to-end distance and the radius of gyration of jR2R3 and jR2R3-P301L. C) Contact map of jR2R3 and jR2R3-P301L. For each frame of the REMD trajectory we considered two residues to be in contact if the minimum distance between any two atoms in the residue were less than 3Å. The color bar represents the fraction of the trajectory in which the two residues were in contact.

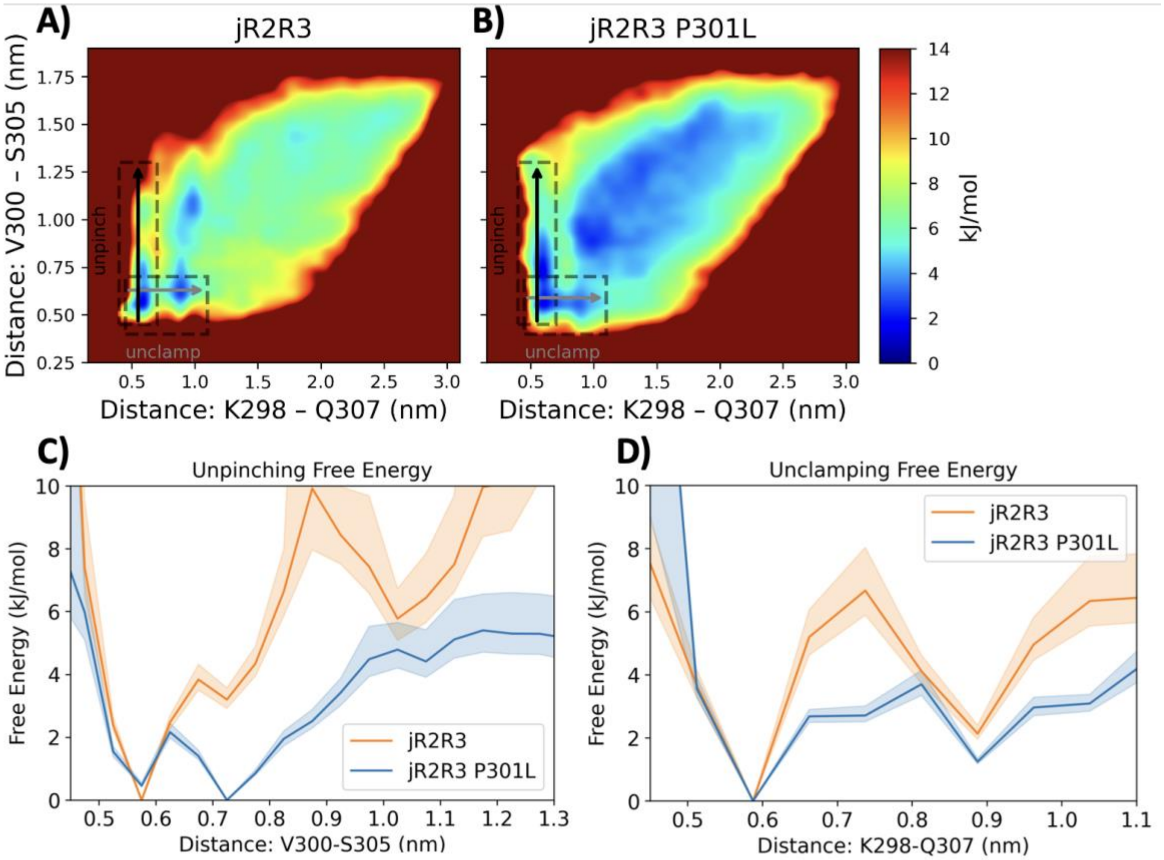


**Fig. S7.** A,B) Pairwise maps described in Figure 4A,B. Boxed regions indicate area indicate are integrated for calculation of panel C and D. C,D) The estimate free-energy of conformational for jR2R3 and jR2R3-P301L in the unpinch (C) and unclamp (D) modes.

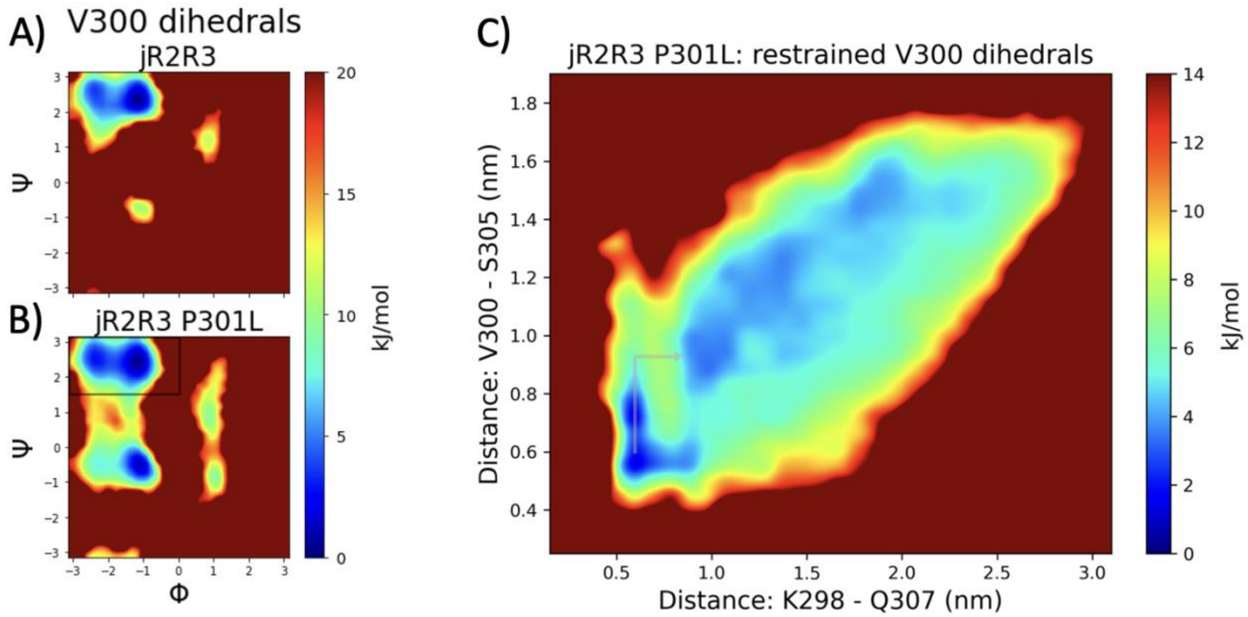


**Fig. S8.** V300 dihedral free energies for A) jR2R3 and B) jR2R3-P301L. The proceeding proline in jR2R3 constrains V300's dihedrals. C) jR2R3-P301L's free energy landscape when only the conformations in the top left of the V300 dihedral free energy landscape are considered. The restricted dihedrals block an unfolding mode (see arrows) where the V300-S305 contact first gets "unpinched" and then the K298-Q307 contact is "unclamped". This unfolding mode is depicted in the supplementary gif.

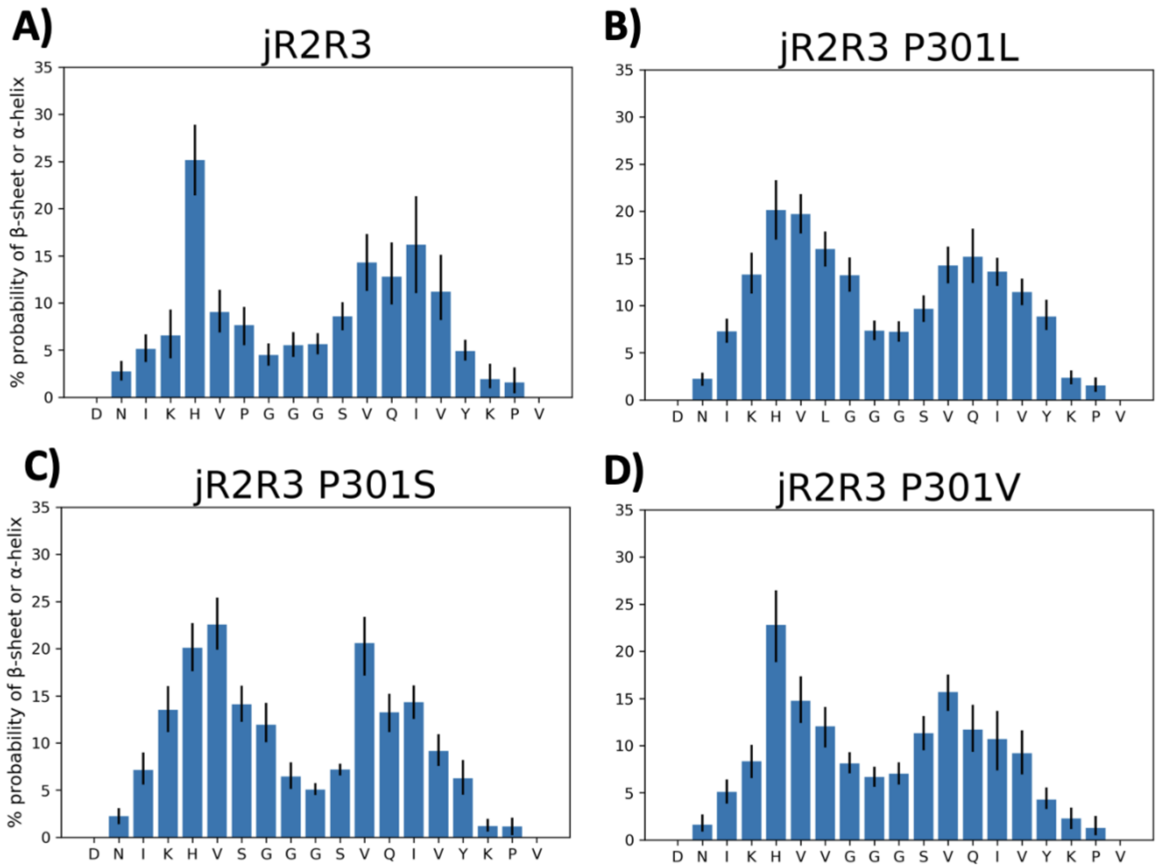


**Fig. S9.** Hydrogen bonding propensity of (A) jR2R3, (B) jR2R3 P301L, (C) jR2R3 P301S, and (D) jR2R3 P301V is measured by β-sheet or α-helix assignment with the DSSP algorithm. 90% confidence error bars are shown. Proline lacks a hydrogen bond donor – the amide nitrogen – so it is generally less prone to forming α-helices and β-sheets.

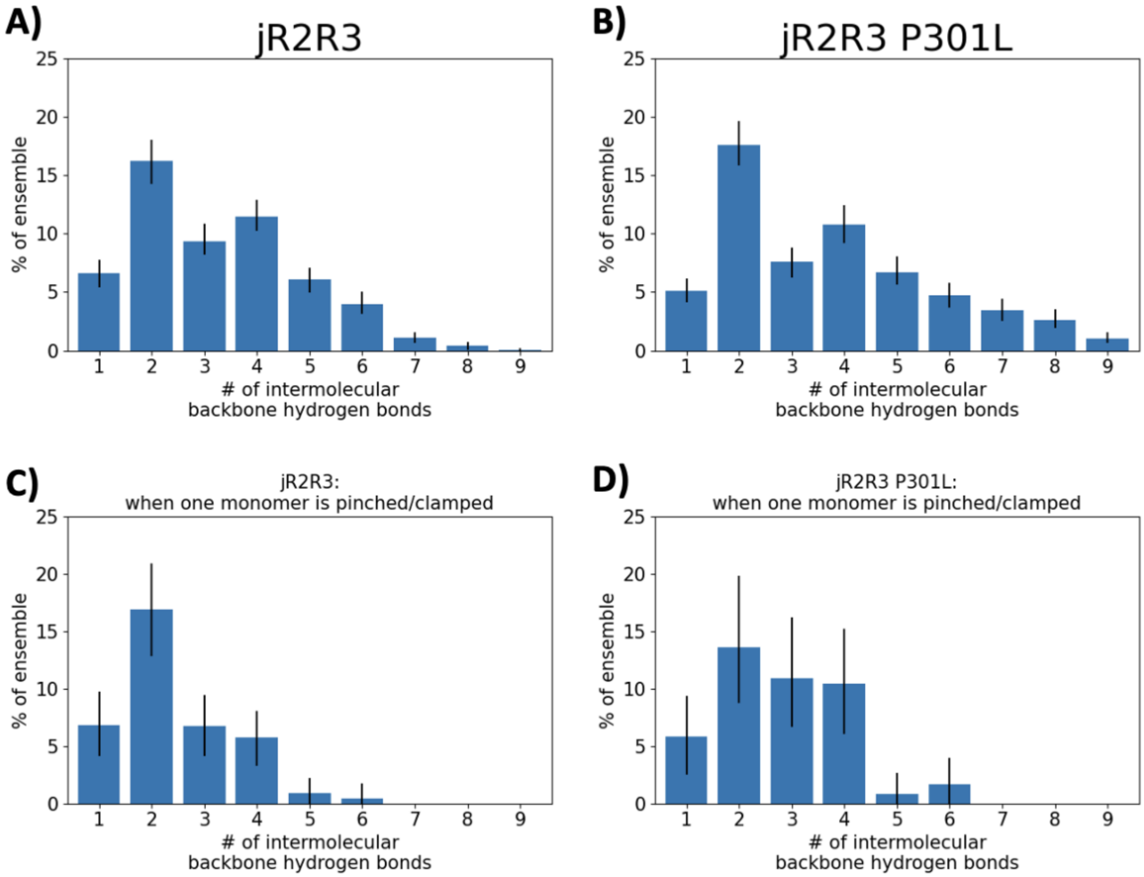


**Fig. S10**. Intermolecular backbone hydrogen bonds for A) jR2R3 and B) jR2R3-P301L dimer simulations. C) and D) show jR2R3 and jR2R3-P301L hydrogen bond probabilities when one of the monomers is closed in a hairpin, i.e. K298-Q307 α-carbon distances < 7Å and V300-S305 α-carbon distances < 8Å. 90% confidence error bars are shown.

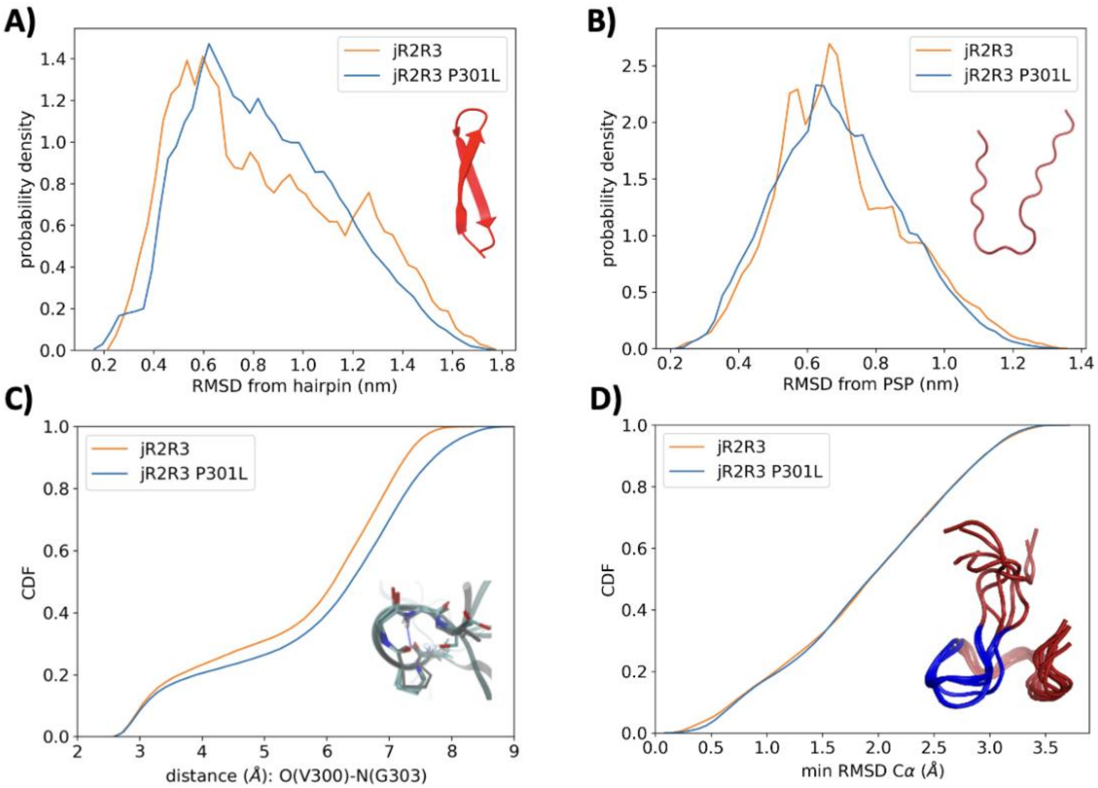


**Fig. S11**. Differences with other simulations of this section of tau in the literature. We observe that wildtype and P301L conformations do not differ to a large extent, contrary to what other researchers have seen with different force fields(37, 38). We use the force field a99SB-disp(39) with TIP4P-D water while Chen, et. al. used a99SB-ILDN with SPCE water and Stelzl, et. al. used a99SB*-ILDN-q with TIP3P water. (A-B) We observe only minor differences between the wildtype and P301 in their RMSD to various hairpins while Chen, et. al. saw significant differences albeit for symmetrical trimers instead of monomers. (A) RMSD distribution with reference to the “best hairpin”, which is the conformation with the highest number of β-sheet residues as determined by the DSSP algorithm. (B) RMSD distribution with reference to the encapsulated hairpin in the PSP disease fold (PDB 7P65, residues 295-313). (C) Cumulative distribution function (CDF) of the distance between O(V300)-N(G303) which interact via hydrogen bonding; this hydrogen bond is depicted with a dashed line. Stelzl, et. al. saw ~5% and ~20% of P301L and WT conformations respectively with an O(V300)-N(G303) distance below 4Å while we see 22-23% for both P301L and WT. (D) CDF of the minimum Cα RMSD of $^{300}$VPGGG$^{304}$ to the closest representative of the NMR ensemble of microtubule-bound tau structures(40) (PDB 2MZ7). Stelzl, et. al. observed ~4% and ~15% of P301L and WT conformations respectively within 1Å of one of the 20 structures in the NMR microtubule-bound ensemble while we see 17-18% for both P301L and WT. Six of the 20 microtubule-bound structures from are depicted with $^{300}$VPGGG$^{304}$ shown in blue.

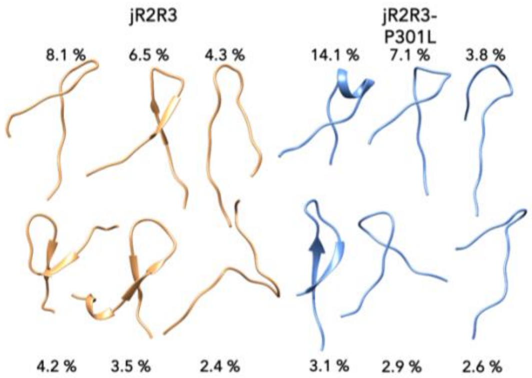


**Figure S12:** Clusters used for INDUS measurements. These clusters were obtained by using the Daura algorithm on monomer REMD simulations.

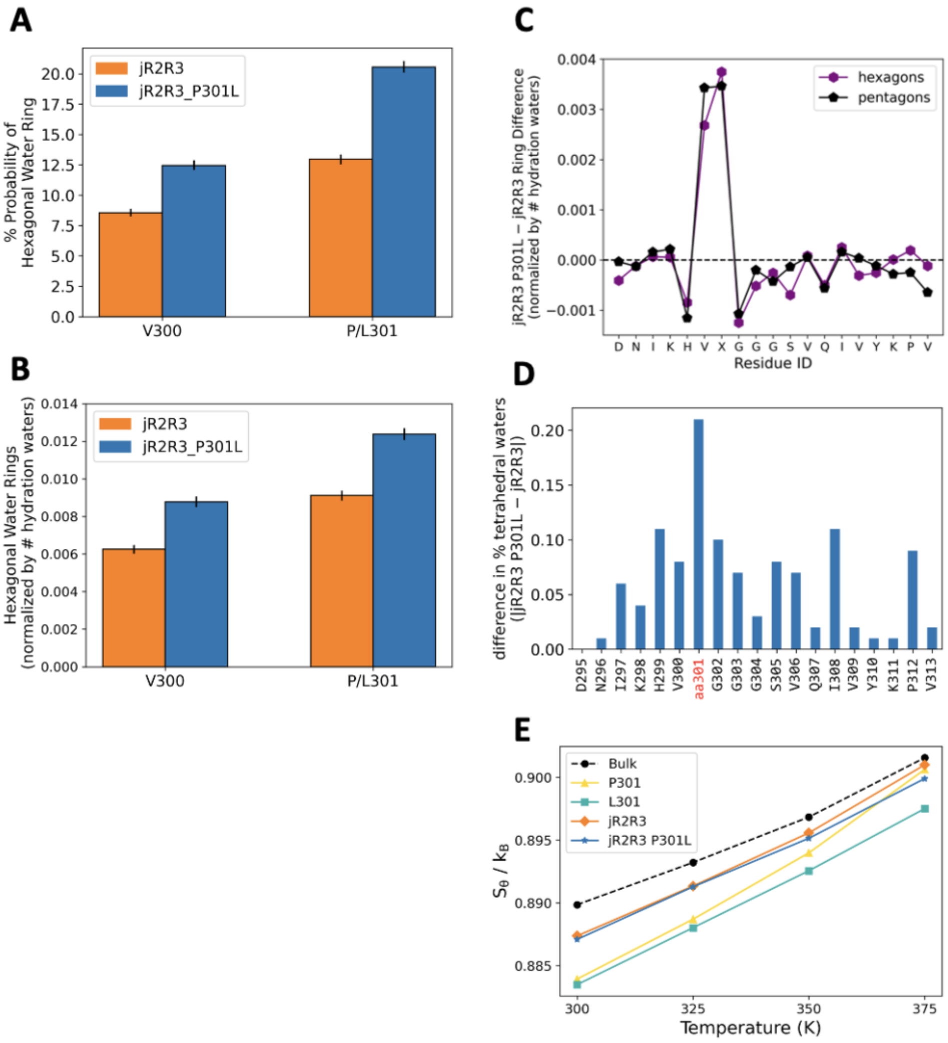


**Fig. S13**. A) the percent probability of surrounding waters having hexagonal character surrounding the V300 and P301(L) residues. B) Difference in relative tetrahedrality (i.e. tetrahedral water fraction divided by the bulk tetrahedral water fraction) for V300's backbone and P/L301's sidechain; 90% confidence error bars are shown. C) Difference in the number or hexagonal and pentagonal water rings formed within 4.25 Å of each residue's heavy atoms; jR2R3 P301L has notably more hexagonal and pentagonal rings formed at V300 and P/L301. D) Difference in jR2R3 and jR2R3-P301L's tetrahedral water fraction around each residue's backbone and sidechain. V300's backbone and P/L301's sidechain show the largest difference in water tetrahedral fraction. E) Shannon entropy ($S_\theta$) measurements of water's 3-body angle distribution for bulk water, around P301 & L301 residues, and around jR2R3 & jR2R3-P301L peptides with varying temperature. This Shannon entropy is correlated with the thermodynamic *excess entropy* (Monroe & Shell, 2019). P301 shows a larger increase in Shannon entropy with temperature than L301.

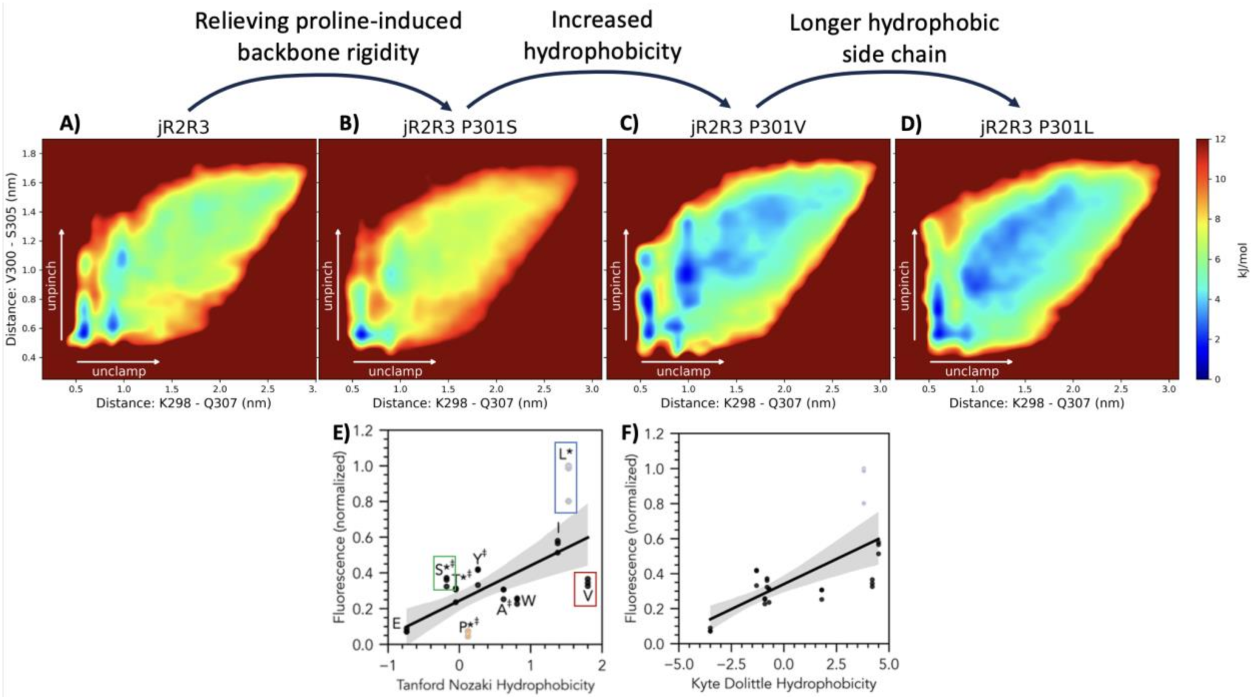


**Fig. S14.** Backbone rigidity, residue hydrophobicity and sidechain length all contribute to the ease of extension for jR2R3. Pairwise energy landscapes as described in Figure 4A,B for (A) jR2R3, (B) jR2R3(P301S), C) jR2R3(P301V), (D) jR2R3-P301L. All the P301 mutants have an energetically accessible unpinch-then-unclamp mode, unlike jR2R3. The free energy wells become less deep with hydrophobicity and hydrophobic length. P301L has a longer hydrophobic side chain than P301V that seems to stabilize intermediates along the unfolding pathway. For each peptide, we conducted REMD simulations of both the HID and HIE protonation states of H299. We show the landscape that most easily opens: HIE for jR2R3 and jR2R3-P301L; HID for P301V; and 50% HID & 50% HIE (Boltzmann-weighted) for P301S. E,F) ThT fluorescence of jR2R3-P301X peptides plotted against different hydropathy scales. (E) Tanford-Nozaki and (F) Kyte-Dolittle.

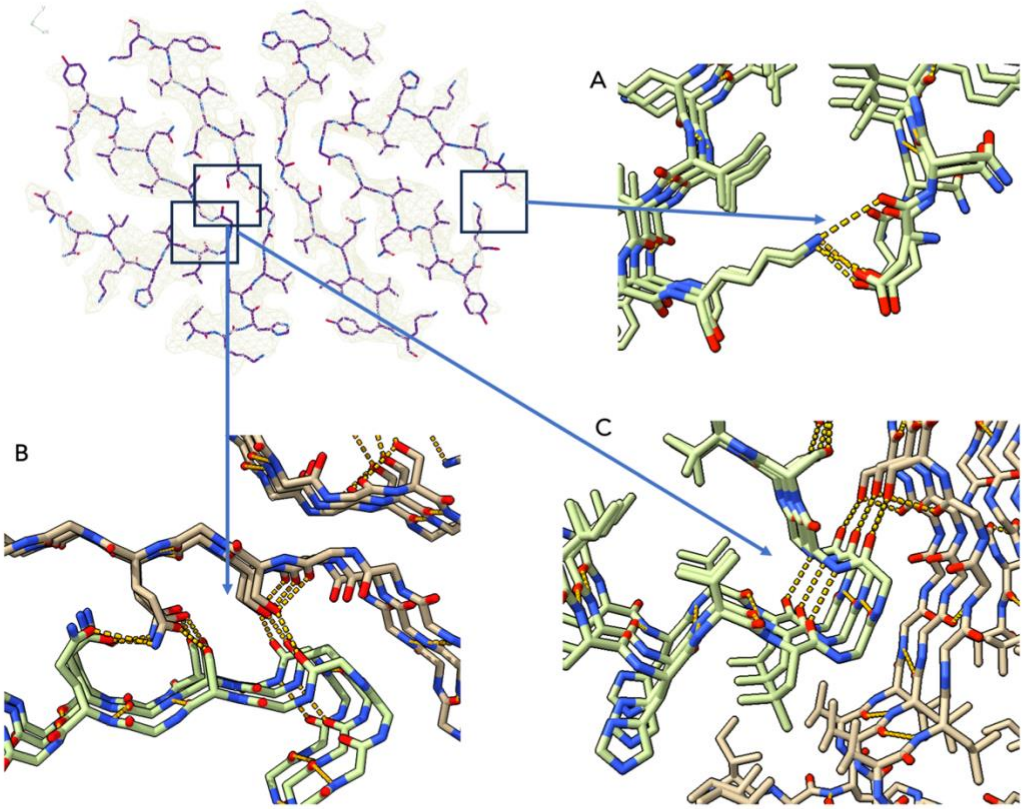


**Fig. S15.** Stabilizing features of the jR2R3-P301L fibrils. Each panel highlights the respective features from the boxed inset. Hydrogen bonds are represented in orange. **A)** The SLS fold (green) is held together at the extremities by an interaction between D295 and K311. **B)** The SLS fold interface with the inner strand (brown) is formed by a three residue bridge between S305 of the SLS strand, Q307 of the inner strand, and Q307 of the SLS strand. S305 of the inner strand may also be forming a hydrogen bond network with G303 of the SLS strand. **C)** the GGG loop of the SLS strand may be held together by a hydrogen bond between the carbonyl of P301L and the amine of G304.

**Table S1.** Cryo-EM data collection, reconstruction, and validation statistics.

| Data Collection | jR2R3-P301L |
|---|---|
| Magnification | x81,000 |
| Defocus Range | -1.0 to -2.0 |
| Voltage (kV) | 300 |
| Microscope | Krios |
| Camera | Falcon 3 |
| Frame exposure time (s) | 3.129 |
| # Movie frames | 50 |
| Total electron dose (e-/Å^2) | 50 |
| Pixel size (Å) | 1.064 |

| Reconstruction | jR2R3-P301L |
|---|---|
| Box size (pixel) | 368 |
| Inter-box distance (Å) | 14.25 |
| # segments extracted | 108,284 |
| # segments after 2D | 85,235 |
| # segments after class3D | N/A |
| Resolution (Å) | 2.98 |
| Map-Sharpening B-factor (Å^2) | -84 |
| Helical Rise (Å) | 2.39 |
| Helical twist (°) | 179.43 |

| Atomic Model | jR2R3-P301L |
|---|---|
| # unique non-hydrogen Atoms | 1440 |
| RMSD bonds (Å) | 0.004 |
| RMSD angles (°) | 0.758 |
| MolProbity score | 2.29 |
| MolProbity clash score | 14.7 |
| Ramachandran favored rotamers (%) | 96.43 |
| Rotamer Outliers (%) | 0 |
| Allowed Rotamers (%) | 3.57 |
| Cβ deviations | 0 |
| Bond outliers (#) | 0 |
| Angle outliers (#) | 0 |
| Correlation Coefficient (Masked) | 0.69 |
